# The metastasis susceptibility gene RRP1B is a stress-responsive regulator of nuclear RNA processing and splicing in breast cancer

**DOI:** 10.64898/2026.06.16.732690

**Authors:** Wan-Ning Li, Ruhul Amin, Thorkell Andresson, Jennifer Steinberg, Michael J. Kruhlak, R. Mark Simpson, Maxwell P. Lee, Kent W. Hunter

## Abstract

The tumor microenvironment exposes cancer cells to mechanical, thermal, hypoxic, and acidic stresses, yet how cells integrate these signals to remodel RNA processing remains poorly understood. Here, we show that ribosomal RNA Processing 1B (RRP1B), previously characterized as a nucleolar ribosome biogenesis factor and metastasis modifier, functions as a broad-spectrum stress sensor that dynamically repositions among the nuclear envelope, nucleolus, and nuclear speckles (NS). Relocalization is governed by multi-site phosphorylation within intrinsically disordered regions (IDRs): phosphomimetic substitutions promote NS-proximal condensate formation in an RNA-dependent manner, while unphosphorylatable substitutions confine RRP1B to the nucleolus. Under stress, the RRP1B interactome shifts globally, with ribosomal processing partners replaced by pre-mRNA splicing components enriched for NS-resident proteins. RNA immunoprecipitation sequencing (RIP-seq) demonstrates that under basal conditions RRP1B associates with long, intron-rich, nuclear periphery-proximal transcripts, whereas heat shock redirects binding toward shorter, exon-dense transcripts enriched for motifs of serine/arginine-rich (SR) proteins near the NS. RRP1B overexpression nearly abolishes cytoplasmic retained intron accumulation and drives preferential export of specific transcript isoforms in a compartment- and temperature-dependent manner, establishing RRP1B as a regulator of RNA localization fidelity rather than transcriptional output. An RRP1B overexpression signature is most highly activated in basal-like and claudin-low breast tumors, and the RRP1B-associated retained intron splicing program correlates with reduced survival in a tumor-grade-dependent manner. These findings reframe RRP1B as a microenvironmentally sensitive regulator of nuclear RNA processing with direct implications for aggressive breast cancer biology.

## Introduction

Nuclear speckles (NSs) are membraneless organelles within the nucleus of eukaryotic cells that are involved in RNA processing^1–3^. These organelles, often found as dozens of foci within a single cell, are rich in splicing regulators and RNA Polymerase II (Pol II)-transcribed transcripts^1^. While this suggests a role in active transcription and RNA splicing, recent evidence shows that NSs are also involved in sophisticated post-transcriptional processes that help cells respond to environmental challenges like stress^4,5^. This expanded role is supported by the discovery that NSs contain not only nascently transcribed RNAs but also transcripts with retained introns (RIs) that can reside stably within speckles^6,7^. It has been proposed that NSs serve as reservoirs for these incompletely spliced transcripts, which are crucial for cell survival. This allows for rapid facilitation of the splicing process on RIs, as these transcripts are in close proximity to splicing factors when cells are under stress. However, the precise mechanism behind this process has yet to be fully defined.

While intron retention is not a new observation^8^, the role of RI-containing RNAs has long been misunderstood, partly due to the assumption that introns are unnecessary for cell growth. In fact, deleting introns in yeast cells significantly impairs their ability to withstand stress, highlighting the essential role of introns in regulating cell survival^9,10^. More recent evidence indicates that RI is not a random mis-splicing event but a form of alternative splicing that affects over 80% of human protein-coding genes.^11,12^. For example, RIs can introduce an additional sequence into an mRNA, potentially leading to premature termination codons and subsequent nonsense-mediated RNA decay^13^. Another subclass of RI transcripts, called detained introns, are predominantly localized in the nucleus and are not spliced or exported to the cytoplasm until external stimuli are encountered^14–17^. These detained introns have been implicated as an adaptive mechanism to stress^18^, though this is still being clarified. A recent study found that, compared to RIs near the inner nuclear membrane, RI-containing transcripts proximal to nuclear speckles tend to have a shorter length and a high GC-content^7^. This suggests that speckle-enriched RIs may be a specific group that is regularly stored in NSs and functionally related to the cellular stress response.

In this study, we demonstrated a previously unrecognized nuclear speckle protein, ribosomal RNA processing 1B (RRP1B), as a key regulator of the post-transcriptional processing of RI transcripts in breast cancer cells under stress conditions. Under standard tissue culture conditions, RRP1B primarily localizes to the granular compartment of the nucleolus, with a small fraction in the nuclear envelope, and has been implicated in the maturation of the large ribosomal subunit^19^. Genetic studies have also demonstrated that polymorphisms of RRP1B predispose breast cancer patients to progress to metastatic disease^20,21^, although the mechanism of action in metastatic progression is currently unknown. Whereas under conditions more representative of physiological tumor microenvironments (physiological stiffness, thermal stress, hypoxia, acidosis), RRP1B was resident in structures other than the nucleolus, most commonly the nuclear speckles. Our data indicate that post-translational regulation (PTM) via serine phosphorylation drives the localization of RRP1B adjacent to nuclear speckles, consistent with heterotypic repulsions underlying microphase formation previously described by Shinn et al.^22^ for speckle-associated intrinsically disordered proteins (IDPs). RNA immunoprecipitation studies further revealed that RRP1B interacts with a broad set of mRNAs, including those containing retained introns. These transcripts are highly enriched for splicing and mRNA export factors, suggesting a role for RRP1B in previously described RNA splicing positive feedback loops^23^. Notably, cellular stress or ectopic expression of RRP1B drives the cytoplasmic accumulation of fully spliced isoforms derived from RRP1B-associated nuclear transcripts undergoing retained-intron splicing. This shift indicates enhanced splicing completion and mRNA export under stress. Importantly, the RI gene signature derived from splicing analysis in RRP1B-bound transcripts is strongly associated with poorer survival in breast cancer patients, linking RRP1B-mediated retained-intron processing to cancer progression.

## Materials & Methods

### Cell lines, culturing, and transfection

Human breast cancer cell lines, MDA-MB-231 and HS578T, were kindly provided by Dr. Jeffrey E. Green (NCI, NIH). Mouse mammary tumor cell lines, 4T1, 4T07, and 67NR were obtained from Dr. Lalage Wakefield (NCI, NIH). HEK293T and HEK293FT cells were purchased from Thermo Fisher Scientific (Invitrogen #R70007). HeLa cells were obtained as a gift from Dr. Pei-Wen Chen from the laboratory of Dr. Paul Randazzo (National Cancer Institute, Bethesda, MD). These cell lines were maintained in Dulbecco’s modified Eagle’s medium (DMEM) (Gibco; high glucose, pyruvate, no glutamine; #10313021) supplemented with 10% fetal bovine serum (FBS) (Gemini SKU#100-106), 1% L-glutamine (Gibco #25030164), and 1% penicillin–streptomycin (Gemini SKU#400-109-100). MCF10A were purchased from ATCC (ATCC # CRL-10317) and were maintained in DMEM/F12 (Invitrogen #11330-032) supplemented with 5% horse serum (Gibco #16050-122), 20 ng/mL EGF (Gibco #AF-100-15), 10 µg/mL insulin (Millipore Sigma #I-1882), 500 µg/mL hydrocortisone (Millipore Sigma #H-0888), 100 ng/mL cholera toxin (Millipore Sigma #C-8052), and 1% penicillin–streptomycin (GiminiBio #400-109).

For liposome transfection, cells were grown on culture dishes 16-18 h beforehand, and transfection was performed using X-tremeGENE™ 9 DNA transfection reagent (Roche #6365779001). The transfection medium was replaced with fresh medium after overnight incubation. After 24 h of incubation, cells were harvested or fixed for experiments. For electroporation, cells were trypsinized and washed once with PBS. After removing residual PBS, cells were mixed with DNA and 100 µL Nucleofector® Solution V (supplement added) (Lonza #VCA-1003), followed by electroporation using program T-018 (Nucleofector® I/II/2b System). Cells were harvested or fixed for experiments after 24 h.

### Plasmids and cloning

#### Gateway Cloning

The complementary DNA (cDNA) of full-length human RRP1B was generated by polymerase chain reaction (PCR) using KOD Hot Start DNA Polymerase (Millipore Sigma #71086-3) (Please find primer sequence in Table S1) and cloned into the Gateway entry vector pENTR1A using the Quick Ligation Kit (NEB #M2200S). The pENTR1A-cDNA entry vector was then used to transfer the encoding region into the lentiviral destination vector pDEST-658 (received as a gift from Dr. Dominic Esposito, NIH) along with a human Pol2 promoter entry vector and N-terminal tag entry vector (GFP or Flag) using the Gateway LR clonase reaction (Thermo Fisher Scientific #11791019).

#### Site-directed mutagenesis for the point mutants

Cloned pDEST658-wild-type human RRP1B vector (with GFP tag) was used to construct vectors for site-directed mutagenesis to generate phosphomimetic mutant (D) and unphosphorylatable mimic (A) for S245, S350, and S702. QuikChange II XL Site-Directed Mutagenesis Kit (Agilent Technologies #200521) was used to perform site-directed mutagenesis according to the manufacturer’s instructions. The integrity of the final constructs was verified through Nanopore whole plasmid sequencing. (Please find primer sequence in Table S1)

#### Combined site-directed mutagenesis and restriction enzyme (RE) cloning for S706 point mutants

Cloned pDEST658-wild-type human RRP1B vector (with GFP tag) was used. PCR fragments containing phosphomimetic mutant (D) and the appropriately close restriction enzyme (RE) digestion sites (BmgBI and AscI) on both ends of the S706 (wild-type DNA sequence: AGT) were individually generated using designed cloning primers (Table S1) by KOD Hot Start DNA Polymerase (Millipore Sigma #71086-3). The PCR reactions were run on 3% 1X TAE agarose, and fragments of the correct size were cut and extracted using QIAquick® Gel Extraction Kit (Qiagen #28704). Two individually purified PCR fragments were mixed and used as the template for the second round of PCR to generate a single amplicon containing phosphomimetic mutant (D) and RE digestion sites (BmgBI and AscI). The PCR reaction was purified by running it on 3% 1X TAE agarose, followed by extraction using the QIAquick® Gel Extraction Kit (Qiagen #28704). Purified PCR fragments and pDEST658-wild-type human RRP1B vector (with GFP tag) were then digested by BmgBI (NEB #R0628; blunt end) and AscI (NEB #R0558; sticky end) according to the manufacturer’s instructions. Fragments of the correct size from each reaction were confirmed and further purified by electrophoresis on 3% 1X TAE agarose and gel extraction using QIAquick® Gel Extraction Kit (Qiagen #28704). Ligation reaction was performed at 16°C overnight using T4 DNA Ligase (NEB# M0202). One Shot™ TOP10 Chemically Competent E. coli (Invitrogen #C404010) was used for transformation according to the manufacturer’s instructions. The same process was used to generate unphosphorylatable mimic (A) in S706. The integrity of the final constructs was verified through Nanopore whole plasmid sequencing. (Please find primer sequence in Table S1)

#### Gibson Cloning for tetracycline (Tet)-inducible S702 mutant

Cloned pDEST658-human RRP1B S702D and S702A mutant vectors (with GFP tag) were used to construct vectors for Tet-inducible S702D and S702A system. Gibson cloning was performed by assembling purified PCR fragments from S702 mutant expression vectors and pCW57-MCS1-P2A-MCS2 (Addgene #80921) using NEBuilder® HiFi DNA Assembly cloning kit (NEB #E5520S), according to the manufacturer’s instructions. The integrity of the final constructs was verified through whole plasmid sequencing. (Please find primer sequence in Table S1)

#### Cloning bimolecular fluorescence complementation (BiFC) vectors

Gateway Entry clones with or without stop codons were used to generate BiFC mammalian expression clones using Gateway™ LR Clonase™ II Enzyme mix (Invitrogen #11791020). Venus BiFC vectors were created by insertion of Venus fluorescent protein coding regions into a Gateway converted pcDNA3.1-zeo vector. Inserted fragments consisted of the N-terminal portion of Venus (amino acids 1-157; VN) or the C-terminal portion of Venus (amino acids 158-238; VC). All vectors were converted to Gateway Destination vectors (Invitrogen #11791020) and subcloned by Gateway LR recombination using the manufacturer’s protocols (Invitrogen #11791020). The RE cloning technique was applied to clone both RRP1B sub-deletions and the point mutants from GFP-tagged vectors (phosphomimetic or unphosphorylatable mimic mutants for S245, S350, and S702) described in the site-directed mutagenesis sections into BiFC vectors.

### Stable cell lines generated using lentiviral vectors

Overexpression stable cell lines, Flag-RRP1B, were made using cloned vectors described in the *Plasmids and cloning* section. For lentivirus production, plasmids and packaging plasmids, psPAX2 (Addgene #12260) and envelope plasmid pMD2.G (Addgene #12259) (obtained from the Trono lab), were transfected into the HEK293FT cell line using X-tremeGENE 9 DNA transfection reagent (Roche; #6365779001). After 48 h of transfection, culture supernatant containing lentivirus was harvested, filtered through a 0.45 μm filter (Millipore #HAWP04700), and then used for transduction of human breast cancer cell lines. Stable cells were selected with 10 μg/mL Blasticidin (Gibco #R21001).

### Treatments for cellular stresses

Cells were grown on tissue culture dishes or coverslips for 24 h with 70-80% confluency before treatments. For heat shock experiments, cells were treated with either 42°C or 37°C for 3 h before harvesting. For hypoxia experiments, cells were treated with either 1% or 21% oxygen for 3 h. For acute acidosis, cells were incubated with media of either pH 6.5 adjusted by HCl or pH 7.4 for 11. h. For mechanical stress, cells were seeded on either hydrogels of different stiffness (Matrigen), including 0.2 kPa and 12 kPa, or glass coverslips (>GPa), coated with Collagen I (50 µg/mL in 20 mM glacial acetic acid) or fibronectin (10 µg/mL in PBS), and harvested or fixed after 48 h.

### Nuclear protein Immunoprecipitation (IP)

Nuclear protein IP was performed using the Nuclear Complex Co-IP Kit (Active Motif #54001) according to the manufacturer’s instructions. Briefly, nuclear lysates were harvested from 15 cm tissue culture dishes and quantified using Pierce BCA Protein Assay Kit (Thermo Fisher Scientific #23225). Subsequently, 200–500 μg of nuclear lysates were incubated with 2 μg of RRP1B antibody (Invitrogen #PA5-53661) or normal rabbit IgG (Cell Signaling Technology #2729) on a rotator at 4 °C overnight. The next day, the reactions were added with 50 μg of Dynabeads Protein G (Invitrogen #10003D) and incubated at 4 °C for 1 h on a rotator. The immune complexes were then washed using a magnetic stand for purification. Finally, the protein condensates were collected from beads by resuspending with 2x reducing buffer (diluted from Invitrogen #NP0009) and eluted after 95 °C incubation for 5 min. (Please refer to Table S1 for more information on the antibodies used in this paper.)

### Liquid chromatography-mass spectrometry (LC-MS)

For RRP1B protein interaction analysis, nuclear protein IP samples, including RRP1B and normal rabbit IgG pulldown, were used and subjected to LC-MS. The experiment was performed by Q Exactive HF (Thermo Fisher Scientific) in the Center for Cancer Research (CCR) Protein Characterization Laboratory (PCL) at Frederick, Maryland.

### Immunoblot analysis

Whole cell protein lysates were prepared by lysis buffer containing 20 mM Tris-HCl (pH 8.0), 400 mM sodium chloride, 1 mM EGTA, 5 mM EDTA, 10 mM sodium fluoride, 1 mM sodium pyrophosphate, 1% Triton X-100, 10% glycerol, and a protease and phosphatase inhibitor cocktail. Nuclear protein lysates were prepared by Nuclear Complex Co-IP Kit (Active Motif #54001) according to the manufacturer’s instructions. The protein concentration was determined using the Pierce BCA Protein Assay Kit (Thermo Fisher Scientific #23225). 20-30 μg of protein lysates were taken for further immunoblot assay by running on NuPAGE 4-12% Bis-Tris protein gels (Invitrogen #NP0322BOX) with MOPS SDS running buffer (Invitrogen #NP0001). Proteins were then transferred onto a PVDF membrane (Millipore #IPVH07850), followed by 1 h blocking with blocking buffer (1xTBST + 5% non-fat dry milk) at room temperature (RT). The membrane was subsequently incubated with primary antibodies at 4 °C overnight, washed with TBST buffer, incubated with secondary antibodies conjugated with horseradish peroxidase (HRP) at RT for 1 h. Chemiluminescence developed from the membrane using the Amersham ECL Prime Western Blotting Detection Reagent (Cytiva #45-002-401) was detected by Amersham ImageQuant 800 imaging system. (Please refer to Table S1 for more information on the antibodies used in this paper.)

### Silver staining

Silver staining was performed using Pierce™ Silver Stain Kit (Thermo Fisher Scientific #24612) according to the manufacturer’s instructions. Briefly, the gel was washed twice with ultrapure water, followed by two times of incubation with fixing solution at RT for 15 min. Gel was then washed twice in 10% ethanol, and twice in ultrapure water, followed by the incubation of the sensitizer working solution for 1 min. After washing twice in ultrapure water, the gel was stained with silver stain for 5 min and subsequently stained with Silver Stain Developer for around 2-3 min until the protein bands appeared. Stop solution was used to wash out the silver stain developer, and acetic acid was used to keep the gel. The gel image was then taken by the Amersham ImageQuant 800 imaging system.

### Immunofluorescence (IF) staining and confocal microscopy

Cells were grown on coverslips for 24 h with 70-80% confluency before treatments or fixation. Fixation was performed using pre-cold methanol at −20 °C for 3 min, followed by cell permeabilization using 0.5% Triton X-100 in PBS at RT for 5 min on a shaker. Cells were then blocked with blocking buffer (0.45 µm filtered 5% bovine serum Albumin in 1x PBS) for 30 min. Subsequently, cells were incubated with primary antibodies diluted in blocking buffer overnight at 4 °C. After three washes with 1xPBS, the cells were incubated with Alexa Fluor-conjugated secondary antibodies for 1 h at RT. Following three additional washes with 1xPBS, ProLong Glass Antifade with NucBlue Stain (Invitrogen #P36981) was applied to preserve the fluorescence signals from cells on glass microscope slides. Images were acquired using a Zeiss LSM 880 Airyscan confocal microscope equipped with a 63x plan-apochromat (N.A. 1.4) oil immersion objective lens with 0.07 um X-Y pixel size, 0.9 um optical section thickness, and when z-stacks were collected a 0.47 um z-step size. (Please refer to Table S1 for more information on the antibodies used in this paper.)

### BiFC analysis

To perform BiFC analysis, cells were transfected with BiFC expression vectors (electroporation for Hs578T cells; liposome transfection for HeLa cells) and seeded on µ-Slide 8 Well chamber slide (ibidi #80806; 1.5 Polymer; ibiTreat). Cells were fixed after 24 h with pre-cold methanol at −20 °C for 3 min, followed by nuclei staining (Hoechst 33342; Invitrogen #H3570) for 10 min at RT. The emission of Venus/YFP fluorescence was detected using Zeiss LSM 880 Airyscan super-resolution confocal microscope [under the same conditions described in the section of *Immunofluorescence (IF) staining and confocal microscopy*].

### RNase A treatment

RNase A treatment was performed after cell fixation using pre-cold methanol at −20 °C for 3 min and permeabilization using 0.5% Triton X-100 in PBS at RT for 5 min. Cells were treated with 0.1 μg/μL RNase A (Millipore Sigma # R4642) for 1 h at 37°C, followed by three additional washes with 1xPBS before IF staining process.

### IF quantification

IF intensity measurements were obtained using ImageJ2 (Fiji). Regions of interest (ROIs) were defined based on NPM1 and DAPI staining to distinguish nucleolar and non-nucleolar regions within the nucleus for analysis of RRP1B intensity. Nuclear size in BiFC images was quantified based on Hoechst 33342 staining.

### Venus (BiFC) chromocenter depletion analysis

Dense heterochromatin foci (chromocenters) were identified within segmented nuclei using Hoechst 33342 signal intensity and a white top-hat morphological transform to determine chromocenters of nuclear area. BiFC interaction foci were detected within the nuclear mask using Laplacian-of-Gaussian blob detection. Spatial enrichment was assessed using a permutation-based approach (10,000 iterations per nucleus), comparing observed spot overlap with chromocenters against randomly distributed points to determine chromocenter depletion.

### RNA immunoprecipitation (RIP)-seq

RIP-seq was performed using RNA ChIP-IT kit (Active Motif #53024) according to the manufacturer’s instructions. Briefly, cells were grown on 15 cm culture dishes with 70-80% confluency. After 24 h incubation and treatments, cells were cross-linked with 1% formaldehyde at RT for 15 min, and the reaction was stopped by the incubation of 0.125 M glycine solution for 10 min. Cells were then collected by gentle scraping and resuspended in lysis buffer (with RNase inhibitors and protease inhibitors) on ice for 30 min after pelleting by centrifugation at 820 rcf for 10 min at 4 °C. The lysates were then centrifuged at 2,400 rcf for 10 min at 4 °C, and the pelleted nuclei were resuspended in Complete Shearing Buffer followed by sonication using Bioruptor. Sheared samples were centrifuged at 18,000 rcf for 10 min at 4 °C, and part of samples (50 µL out of 350 µL in total) were taken for RNA quantification before performing IP. 10 µg of RNA per IP reaction were incubated with specific primary antibodies, along with 25 μL of Dynabeads Protein G magnetic beads (Invitrogen #10-003-D), on a rotator at 4 °C for overnight. The immune complexes were washed with RNA-ChIP Wah Buffer and collected with elution buffer utilizing a magnetic stand to pellet beads. With the addition of 0.1M NaCl and proteinase K, the samples were incubated at 42 °C for 1 h, followed by 65 °C for 1.5 h for reverse crosslinking and protein digestion to release RNA. RNA purification was performed using TriPure Isolation Reagent (Roche #11667157001) as well as RNA Clean & Concentrator-5 (Zymo #R1013). The samples were then treated with DNase I at 22 °C for 25 min, stopped by Stop solution, followed by RNA cleaning utilizing RNA Clean & Concentrator-5 again. The purified RNA was subjected to library preparation and sequencing performed on the Nextseq 2000 system (Illumina) at CCR Sequencing Facility, Frederick, Maryland. (Please refer to Table S1 for more information on the antibodies used in this paper.) The RIP-seq samples generated 23–32 million pass-filter reads, with approximately 90% of bases exceeding a quality score of Q30. Reads were aligned to the Telomere-to-Telomere version 2 (T2T v2) genome for most analyses, except for rMATS and integration with subcellular RNA-seq data, which were performed using the GRCh38 (hg38) genome due to compatibility requirements.

### RIP-seq data analyses

#### Replicate Multivariate Analysis of Transcript Splicing (rMATS)

Alternative splicing analysis was performed using Replicate Multivariate Analysis of Transcript Splicing (rMATS) turbo v4.3.0, including skipped exon (SE), retained intron (RI), alternative 3′ splice site (A3SS), alternative 5′ splice site (A5SS), and mutually exclusive exon (MXE) events. Significant splicing events were defined by a false discovery rate (FDR) ≤ 0.05.

#### Intron retention ratio (IRR)

For each annotated intron, reads were counted from sorted BAM files using pysam. Intronic reads were defined as reads whose alignment fell entirely within intron boundaries with no splice junction (no N operation in the CIGAR string). Exonic reads were counted from the flanking exons. IRR was computed per intron per replicate as intronic reads / (intronic reads + exonic reads), then averaged across replicates. The stress-induced change was quantified as ΔIRR = IRR(42°C) − IRR(37°C). Introns were classified as significantly stress-decreased or stress-increased if both replicates showed a consistent direction of change and |ΔIRR| > 0.1.

#### Splice Site Strength

Splice site strength was quantified using the maximum entropy model of Yeo and Burge^24^ as implemented in the MaxEntScan framework. For each unique intron, two splice site windows were extracted. The 5′ splice site (5′SS) window comprised a 9-mer spanning 3 nucleotides of exonic sequence immediately upstream of the intron and 6 nucleotides of intronic sequence beginning at the splice junction (positions −3 to +6 relative to the exon–intron boundary). The 3′ splice site (3′SS) window comprised a 23-mer spanning 20 nucleotides of intronic sequence ending at the splice junction and 3 nucleotides of downstream exonic sequence (positions −20 to +3 relative to the intron–exon boundary). For minus-strand introns, the genomic sequence of the complementary window was extracted and reverse-complemented prior to scoring. Only canonical GT-AG introns, defined as those with GT at positions 4–5 of the 5′SS 9-mer and AG at positions 19–20 of the 3′SS 23-mer, were retained for scoring. Two-sided Mann-Whitney U test with Benjamini-Hochberg FDR correction was used for group comparisons.

#### Splicing index (SI)

Junction reads (reads containing an N operation in the CIGAR string) and intronic reads were counted per gene across all four BAM files (two replicates × two temperatures) using pysam against the T2T-CHM13v2 RefSeq GTF. Replicate counts were summed per condition. SI was computed per gene per condition as junction reads / (junction reads + intronic reads). For the background comparison, SI was computed for 16,033 additional genes not present in the RIP-seq dataset that passed the same read depth filter.

### Subcellular RNA isolation and sequencing

Cells were grown in 15 cm tissue culture dishes with 70-80% confluency. Three replicate dishes were used for each condition. Reagents from the RNA ChIP-IT kit (Active Motif #53024) were used to perform the subcellular fractionation process. Nuclear and cytoplasmic RNA fractions were purified using TriPure Isolation Reagent (Roche #11667157001) and RNA Clean & Concentrator-5 (Zymo #R1013). Briefly, one volume of TriPure Isolation Reagent was added to the nuclear or cytoplasmic fractions for total RNA extraction. Chloroform (0.2 volumes) was then added to the homogenized samples, followed by centrifugation at 12,000 × g for 15 min at 4 °C. The upper aqueous phase (approximately 0.5 volumes), containing the RNA, was transferred to a fresh tube and further purified using the RNA Clean & Concentrator-5 kit according to the manufacturer’s instructions. The samples were then treated with DNase I at 22 °C for 25 min, stopped by Stop solution, followed by RNA cleaning utilizing RNA Clean & Concentrator-5 again. The purified subcellular RNA samples were subjected to library preparation and sequencing performed on NovaSeq Xplus 1.5B (Illumina) at the CCR Sequencing Facility, Frederick, Maryland. The subcellular RNA samples generated 120–190 million pass-filter reads, with approximately 92% of bases exceeding a quality score of Q30. Reads were aligned to the hg38 genome for further analysis.

### Nuclear-to-cytoplasmic (N/C) ratio analysis of Subcellular RNA-seq

N/C ratio analysis performed at the transcript isoform level. Transcripts with a maximum group-mean TPM < 1 across all eight experimental groups were excluded, retaining 65,832 transcripts for analysis. For each transcript, log2(TPM + 1) values were computed and per-replicate N/C ratios calculated as the difference between matched nuclear and cytoplasmic log2-TPM values. Five comparisons were tested: RRP1B overexpression at 37°C, RRP1B overexpression at 42°C, heat shock in parental cells, heat shock in oeRRP1B cells, and an overexpression status-by-temperature interaction term. For each comparison, a one-sample t-test (μ = 0) was applied to the three-replicate delta-N/C values per transcript. Transcripts were considered significantly shifted if nominal p < 0.05 and |Δlog2(N/C)| ≥ 0.5.

### Histology and immunohistochemistry (IHC) staining

The procedure for IHC staining was described previously^25^. Briefly, tissue sections were deparaffinized by immersion in HistoChoice (Amresco, Inc., Solon, OH) for 10 min, rehydrated followed by antigen retrieval with citrate buffer pH 6.1 (Target Retrieval Solution 10X, Agilent Dako # S169984-2). Tissue sections were washed and quenching of endogenous peroxidase was performed by 3% hydrogen peroxide for 5 min. Slides were blocked for 30 min with serum-free protein block (Agilent Dako # X090930-2). After washing, slides were treated for 1 h with the corresponding biotinylated secondary antibody (Thermo Fisher Scientific, 1:200) diluted in 3% BSA solution, followed by avidin-biotinylated conjugation using the ABC system (Vector Labs) for 30 min. ImmPACT DAB (Vector Lab # SK-4105) was used for staining development following the manufacturer’s suggestions. Counterstaining was performed with Hematoxylin QS (Vector Lab # H-3404) for 1 min and water wash. All slides were dehydrated and mounted with VectaMount (Vector Lab # H-5501-60). (Please refer to Table S1 for more information on the antibodies used in this paper.)

### RNA fluorescence in situ hybridization (FISH) combined with IF staining

Cells were grown on coverslips 24 h with 70-80% confluency before treatments or fixation. RNA FISH experiments were performed using the HCR™ Gold RNA-FISH kit (Molecular Instruments) according to the manufacturer’s instructions. RNA probes targeting individual gene transcripts were designed by the Molecular Instruments team. Briefly, fixation was performed using 4% Formaldehyde at RT for 10 min, followed by Pre-hybridization by HCR™ HiFi Probe Hybridization Buffer at 37°C for 30 min. Subsequently, cells were incubated with HCR™ HiFi Probe diluted in Probe Hybridization Buffer overnight at 37 °C. After four washes by HCR™ HiFi Probe Wash Buffer at 37 °C every 15 minutes, the cells were incubated with HCR™ Gold Amplifier Buffer at RT for 30 min. In the meantime, snap-cooled h1 hairpins and snap-cooled h2 hairpins were prepared separately by heating at 95 °C for 90 sec and cooling to RT in the dark for 30 min. Cells were incubated with the 1:1 mixture of snap-cooled h1 and h2 hairpins in HCR™ Gold Amplifier Buffer at RT for 4 h. After four washes by HCR™ Gold Amplifier Wash Buffer at 37 °C every 15 minutes, cells were post-fixed with 4% Formaldehyde at RT for 10 min. Cells were then blocked with blocking buffer (0.45 µm filtered 5% bovine serum Albumin in 1xPBS) for 30 min. Subsequently, cells were incubated with primary antibodies diluted in blocking buffer at RT for 1 h. After three washes with 1xPBS, the cells were incubated with Alexa Fluor-conjugated secondary antibodies for 1 h at RT. Following three additional washes with 1xPBS, ProLong Glass Antifade with NucBlue Stain (Invitrogen #P36981) was applied to preserve the fluorescence signals from cells on glass microscope slides. Images were acquired using a Nikon SoRa Spinning Disk confocal microscope equipped with a 60x CFI Apo TIRF (N.A. 1.49) oil immersion objective lens and Hamamatsu ORCA Fusion BT sCMOS camera with 0.11 um X-Y pixel size and 1.0 um optical section thickness. (Please refer to Table S1 for more information on the antibodies used in this paper.)

### Data availability

Sequence data will be made available through SRA prior to publication of this manuscript.

## Results

### Tumor-associated microenvironmental stresses induce RRP1B re-localization

Recent work from our laboratory demonstrated that RRP1B is responsive to extracellular mechanical stiffness, with increased protein levels when cells were plated on higher tensile strength matrices (Fig. S1A). Unexpectedly, when immunofluorescence assays were performed on cells plated on 0.2 kPa hydrogels, which approximate normal breast tissue stiffness, the majority of RRP1B was localized to the cytoplasm (Fig.1A and Fig. S1B). In contrast, on 12 kPa hydrogels, which approximate tumor stiffness, RRP1B was found primarily in the nucleus and was almost exclusively in the nucleolus on >GPa plastic dishes under standard tissue culture conditions, demonstrating RRP1B responds to extracellular stiffness with changes in both protein levels and subcellular localization. To determine whether these changes were physiologically relevant, immunohistochemistry staining on human patient tissue arrays was performed. Consistent with the *in vitro* analysis, RRP1B was found to be nuclear in both adenocarcinoma and ductal carcinoma *in situ*, while in normal human breast epithelia RRP1B staining was significantly reduced (Fig.1B).

**Figure 1.**
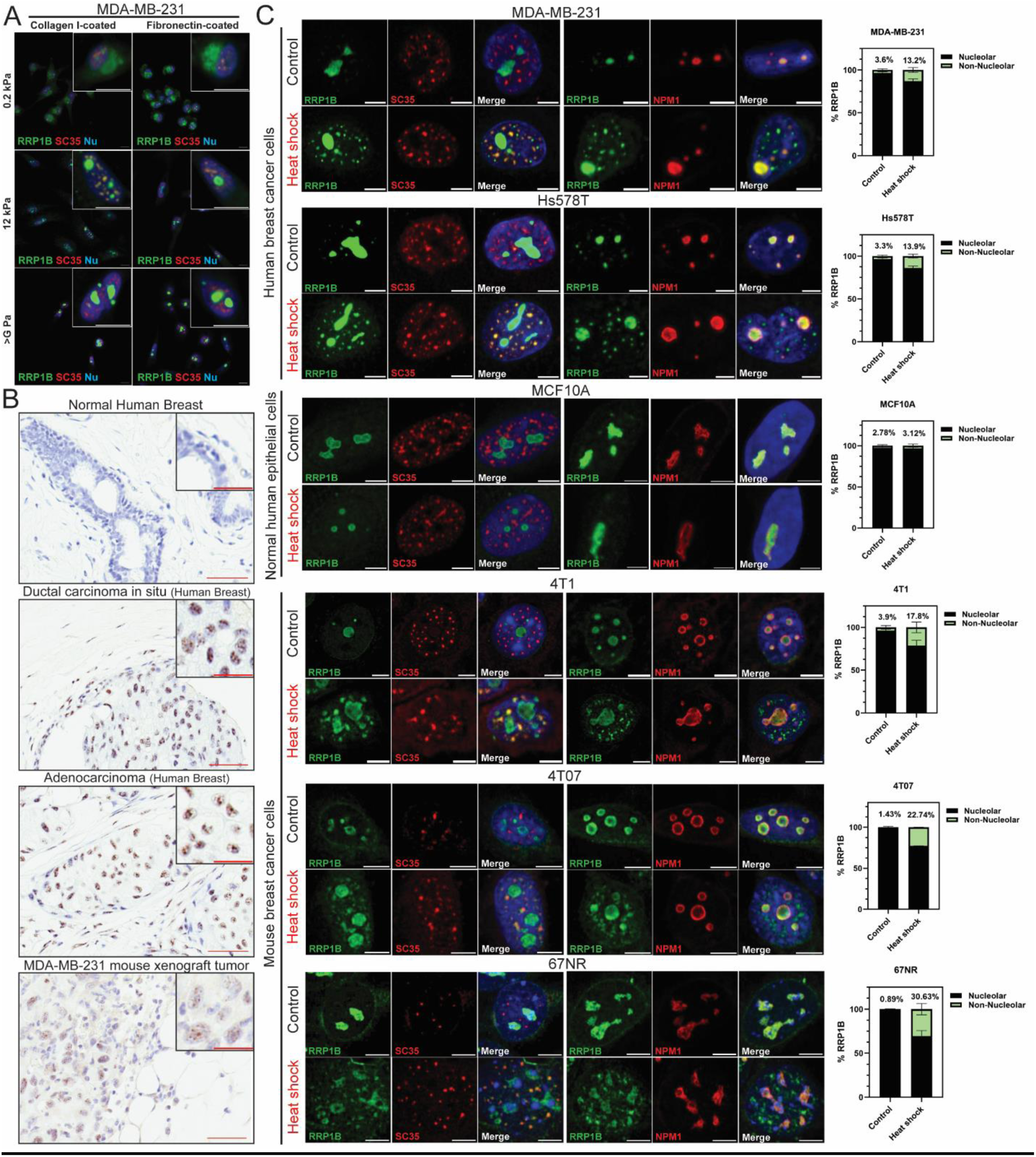
RRP1B protein localization under tumor-associated microenvironmental stresses. (A) Representative immunofluorescence staining confocal images of MDA-MB-231 cells seeded on hydrogels of different stiffness (0.2 kPa, 12 kPa, or >G Pa) that were coated with either collagen or fibronectin. Scale bar, 10 μm. (B) Representative immunohistochemistry staining images of normal human breast, human breast cancer tissues, and mouse xenografts stained with RRP1B antibody using DAB, with nuclei counterstained with hematoxylin. Scale bar, 50 and 25 μm. (C) Representative immunofluorescence staining images and quantitative analysis of human breast cancer cells (MDA-MB-231 and Hs578T), normal human breast epithelial cells (MCF10A), and mouse breast cancer cells (4T1, 4T07, and 67NR) cultured at either 37°C or 42°C for 3 hours. Scale bar, 5 μm. (Two-way ANOVA with Fisher’s LSD test.)

To better understand the microenvironmentally induced response of RRP1B, RRP1B localization was examined in cells subjected to other tumor-relevant environmental stresses, including thermal stress, acidosis, and hypoxia. Immunofluorescence analysis revealed enrichment of RRP1B in extra-nucleolar structures for all tumor-associated environmental stresses (Fig.1C and Fig. S1C-E). A small fraction of RRP1B observed initially localized to the inner face of the nuclear membrane is also lost after heat shock (Fig. 1C and Fig. S1C). Stress-associated re-localization of RRP1B was conserved across mouse mammary tumor and human breast cancer cell lines but was less pronounced in normal human breast epithelial cells (Fig. 1C). This suggests that the stress response is not a cell line– or species-specific artifact, but rather a feature more specific to cancer cells in response to microenvironmental stress. Co-staining with SC35, which targets the SON protein, revealed that the extranucleolar RRP1B co-localized with the membrane-less NS condensate. Fluorescence intensity analysis revealed that the stress-induced extra-nucleolar RRP1B signal increases between ∼4-30-fold, depending on the cell line and stress conditions (Fig. 1C and Fig. S1F-G). Taken together, these data suggest a previously unrecognized response of RRP1B to tumor-associated microenvironmental stresses.

### RRP1B localization is mediated by multiple intrinsically disordered regions (IDR) phosphorylation events

RRP1B is a 758–amino acid protein comprising a single predicted structured Nop52 domain in its N-terminal third, with the majority of the remaining sequence predicted to consist of intrinsically disordered regions (IDRs) (Fig. S2A). IDRs are thought to mediate interactions with proteins and/or nucleic acids through post-translational modifications and/or contact-mediated conformational changes in structure^26–28^. Consistent with this, previous studies have demonstrated that phosphorylation of S702 within the IDR mediates RRP1B interaction with PP1β or PP1γ in the nucleolus^29^. Analysis of the protein sequence (NP_055871.1) using the NLS Mapper^30,31^ and the Nucleolar Localization Sequence Detector^32^ identified four predicted nuclear localization sequences (NLS) and multiple predicted nucleolar localization sequences (NoLS) overlapping the largest IDR (amino acids 381-598), as predicted by MobiDB^33^, within the C-terminal region of the protein (Fig. S2A-C). Deletion of the entire C-terminal region of RRP1B (Full ΔCterm: 1-258 a.a.) led to extranucleolar localization (Fig. 2A). Further deletion studies from our laboratory have also previously suggested that the middle and C-terminal third of RRP1B contribute to the nuclear envelope and nuclear localization, respectively^21^. To further refine the understanding of the molecular basis of RRP1B nuclear speckle localization, deletion and point mutation analysis were performed.

**Figure 2.**
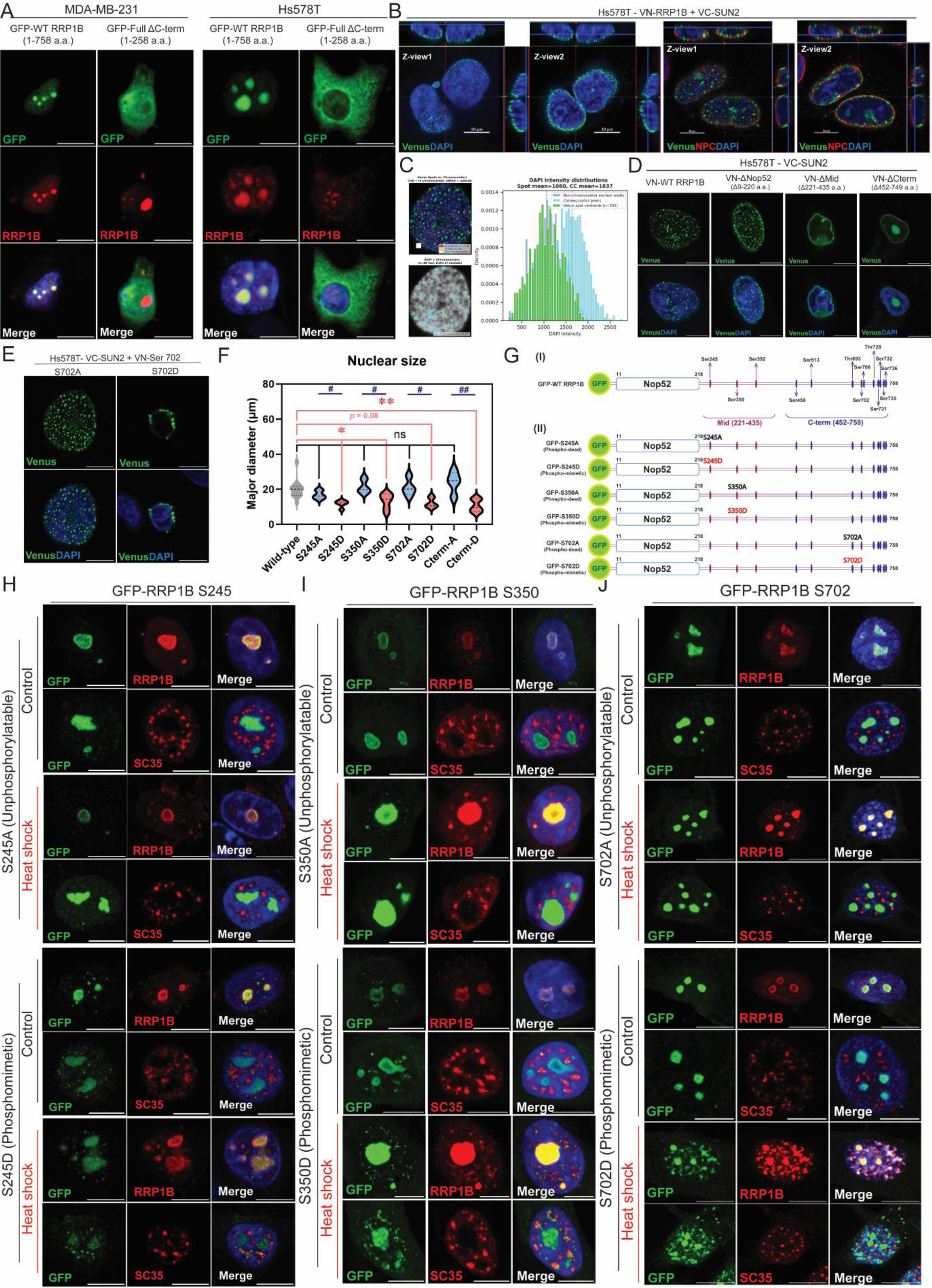
Regulation of RRP1B localization by serine phosphorylation at the C-terminal intrinsically disordered regions. (A) Representative immunofluorescence staining images of MDA-MB-231 and Hs578T cells expressing either GFP-WT RRP1B or GFP-Full ΔC-term proteins. Scale bar, 10 μm. (B) Representative Z-stack BiFC images of Hs578T co-transfected with VC-SUN2 and VN-RRP1B after 24 hours. (NPC: nuclear pore complex. VC or VN: C or N-terminal half of Venus protein tag) (C) Chromocenter image quantitative analysis on BiFC images of Hs578T co-transfected with VC-SUN2 and VN-RRP1B after 24 hours. (D) Representative BiFC images of HS578T cells co-transfected with VC-SUN2 and VN-RRP1B deletion constructs. Scale bar, 10 μm. (E) Representative BiFC images of HS578T cells co-transfected with VC-SUN2 and VN-RRP1B containing either the S702A (unphosphorylatable) or S702D (phosphomimetic) mutation. Scale bar, 10 μm. (F) Quantitative analysis of nuclear size in HS578T cells co-transfected with VC-SUN2 and VN-RRP1B containing either unphosphorylatable or phosphomimetic mutations. (p < 0.05, p < 0.01; one-way ANOVA with Šídák’s multiple-comparisons test. #, p < 0.05, ##, p < 0.01; unpaired t-test.) (G) Schematic representation of the RRP1B point mutation constructs used to determine the residues regulating subnuclear localization. (H-J) Representative immunofluorescence images of HS578T cells expressing GFP-tagged RRP1B proteins with either unphosphorylatable or phosphomimetic mutations cultured at either 37°C or 42°C for 3 hours. Scale bar, 10 μm.

Due to the relatively low concentration of RRP1B at the nuclear envelope in human triple-negative breast cancer (TNBC) cells we employed biomolecular fluorescent complementation assays (BiFC)^34^ to better visualize RRP1B at this location. BiFC utilizes split Venus as epitope tags. When proteins tagged with the N- or C-terminal tags are in close proximity, the Venus halves can refold to generate a fluorescent signal. Transfection of BiFC constructs for RRP1B and the inner nuclear membrane protein, SUN2, into TNBC cells results in a fluorescence punctate pattern in the nuclear membrane (Fig. 2B, left 2 panels). Co-staining with DAPI revealed that the RRP1B/SUN2 nuclear membrane BiFC puncta were depleted for DAPI-rich regions (N= 25 cells, p = 2.4E-32) (Fig. 2C), suggesting these structures are enriched in euchromatic regions near the inner nuclear membrane. Co-staining for the nuclear pore complex further demonstrated that the RRP1B–SUN2 BiFC signal does not co-localize with nuclear pores (Fig. 2B, right two panels), indicating that this structure is independent of the previously described NUP210–SUN2 metastasis-associated nuclear membrane complex^35^. Deletion of Nop52 domain (ΔNop52) did not alter the punctate nuclear membrane signal (Fig. 2D and Fig. S2D). In contrast, deletion of either the middle third of RRP1B (ΔMid) or the C-terminal third (ΔCterm) resulted in diffuse staining of the nuclear envelope along with nucleolar localization (Fig. 2D), indicating that the C-terminal region is essential for proper nuclear membrane localization.

To further determine whether phosphorylation of the PP1β and PP1γ S702 contact residue played a role in RRP1B nuclear envelope localization, RRP1B S702A unphosphorylatable and S702D phosphorylation mimetic mutations were analyzed. RRP1B S702A displayed the wild-type punctate nuclear membrane pattern, consistent with all of the deletion mutants encompassing this region (Fig. 2E). In contrast, the S702D phosphomimetic mutant resulted in fluorescent clustering in large bodies at the nuclear periphery and reduced nuclear size (Fig. 2E-F). To determine if other IDR phosphorylation sites might be necessary for proper RRP1B localization additional phosphomimetic or phosphorylation-dead mutations were generated. Analysis of the PhosphoSitePlus database (https://www.phosphosite.org) identified phosphorylation as the predominant PTM on human RRP1B protein, with 48 reported phosphorylation sites (Fig. S2E). Since the deletion analysis of the middle third of the protein (ΔMid) suggested a potential role for this region in nuclear membrane localization^36^ (Fig. 2D and Fig. S2D), we also analyzed the S245 and S350 phosphorylation sites, given their high representation in reference datasets and conserved sequences across species (Fig. S2E and Supplementary PRALN file). Both the S245A and S350A mutants retained the punctate pattern observed for wild type RRP1B (Fig. S2F). In constrast, the phosphomimetic S245D and S350D showed a mixed pattern, with some cells displaying the wild-type punctate pattern and other cells displaying the large aggregated fluorescence (Fig. S2F). This suggests constitutive phosphorylation of any single site, mimicked by the S-to-D mutations, increases the propensity of the protein to form large aggregates, potentially by phase separation mediated by IDR.

To investigate the localization of nucleoplasmic RRP1B, GFP-tagged expression constructs were generated. Introduction of S245A, S350A, S702A, or S706A point mutants, which are conserved serine sites across species and highly represented in the PhosphoSitePlus database, into HS578T cells resulted in signal almost exclusively within the nucleolus at either control or after heat shock (Fig. 2G-J, Movie 1, and Fig. S2G-H). In contrast, the phosphomimetic mutants (S245D, S350D, S706D) frequently formed small puncta (> 1 mm in diameter), adjacent to, but not integrated into, the SC35-positive nuclear speckle under both conditions (Fig. 2H-I, movie 2, and Fig. S2H). Puncta formation was sensitive to RNase treatment (Fig. S2J), suggesting that these structures may represent microphase condensates^22^ formed upon ectopic expression of RRP1B, which may prevent full integration into nuclear speckles. For the phosphomimetic S702 (S702D), the GFP signals redistributed predominantly to regions proximal to nuclear speckles, which was enhanced under thermal stress (Fig. 2J). Taken together, these data suggest that RRP1B localization at the nuclear speckle is mediated by a set of complex, multi-residue phosphorylation events within its intrinsically disordered domains.

### RRP1B associates with splicing components in response to cellular stress

To study the potential role of RRP1B in the NS under stress, RRP1B immunoprecipitation followed by mass spectrometry (IP-MS) was performed under standard (37 °C) or heat shock (42 °C) conditions (Fig. 3A and Fig. S3A-C). A total of 891 putative RRP1B-interacting candidates were identified. Of these proteins, 136 proteins (15.3%) exhibited increased association with RRP1B under heat shock, 681 proteins (76.4%) showed decreased binding, and 74 proteins (8.3%) showed no change (Fig. 3B and Table S2). Consistent with previously published findings^29^, both PP1β and PP1γ showed decreased interaction with RRP1B under stress conditions (Table S2). Globally, proteins with decreased association with RRP1B were enriched for ribosomal processing (Fig. 3C, upper panel), consistent with known downregulation of these pathways during thermal stress^37,38^ and with previous associations of RRP1B with 60S ribosomal subunit processing and nuclear export^19,23^. In contrast, the most significant ontology associated with proteins exhibiting an increased RRP1B-association after heat shock was processing of capped intron-containing pre-mRNA (REAC:R-HSA-72203 (Fig. 3C lower panel, 3D, and Fig. S3D-E), consistent with increased RRP1B concentrations in the NS and previous associations of RRP1B with splicing. Of the 136 proteins showing increased association with RRP1B after cellular stress, 76 have been previously associated with RNA processing, with 15 previously characterized as nuclear speckle proteins (GO:0016607; p-value: 2.13E-09) (Fig. 3D-E). The changes in binding indicate that RRP1B preferentially associates with nuclear speckle proteins rather than ribosomal or other proteins under stress conditions (Fig. 3F). Intriguingly, these nuclear speckle-associated proteins included Xenopus kinesin-like protein 2 (TPX2), which was previously identified as the central protein of a published metastasis-associated network^39,40^ (Table S2). Together, these data therefore suggest a potential role for RRP1B in modulating both ribosomal and RNA processing in response to cellular environmental stress.

**Figure 3.**
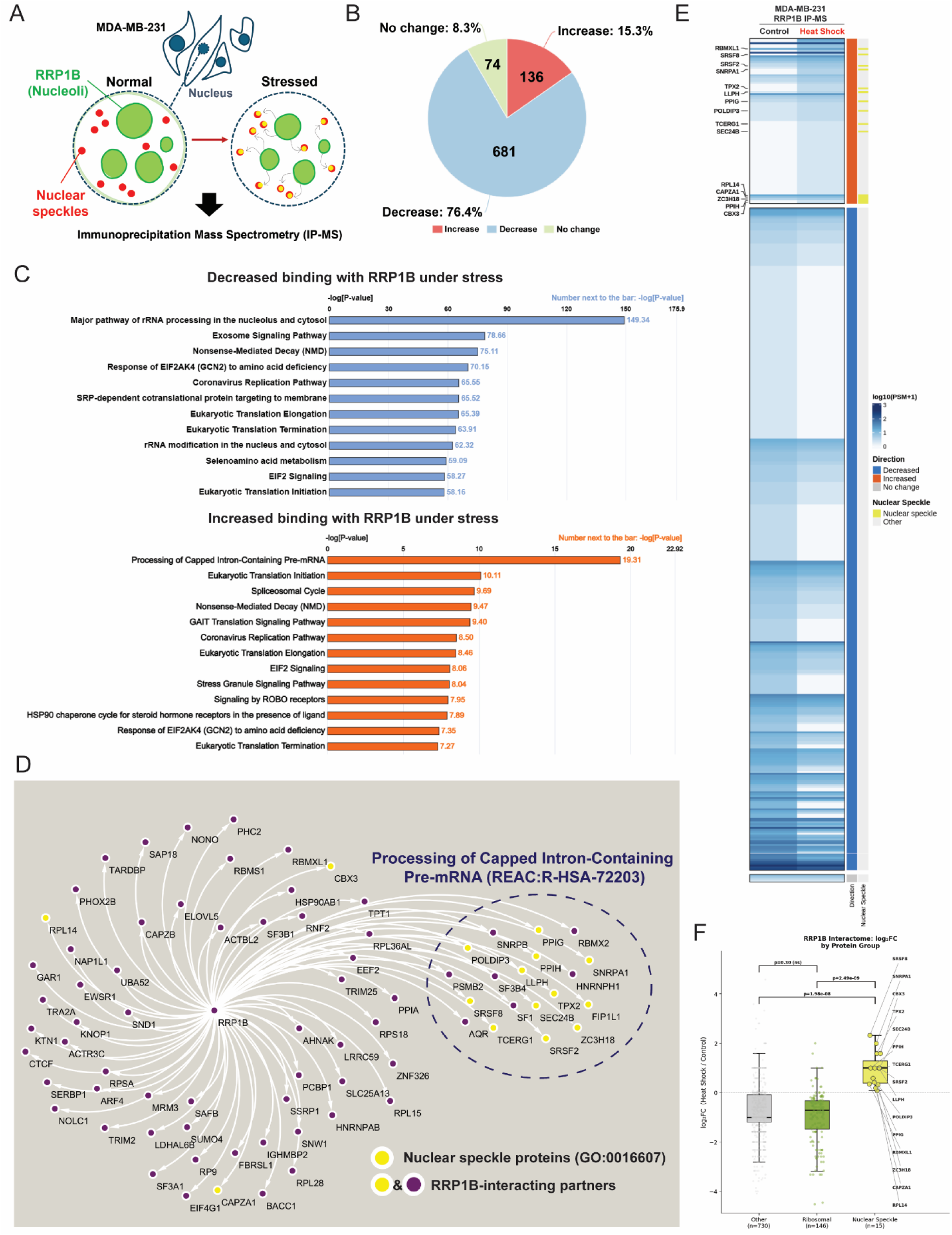
Direct interaction of RRP1B with nuclear speckle proteins in response to cellular stress. (A) Schematic diagram of the RRP1B IP-MS workflow performed in MDA-MB-231 cells. (B) Pie chart illustrating the three major categories of RRP1B-interacting proteins classified by stress-induced changes in peptide-spectrum matches (PSMs) from RRP1B IP-MS analysis. (C) Top canonical pathways identified by Ingenuity Pathway Analysis (IPA) among proteins with decreased (n = 681; upper panel) or increased (n = 136; lower panel) binding to RRP1B under stress conditions. (D) Cytoscape network analysis of 76 proteins with increased binding to RRP1B that are functionally associated with RNA processing. (E) Binding heatmap of RNA processing proteins exhibiting differential association with RRP1B in response to heat shock. (F) Boxplot of fold changes in RRP1B binding under heat shock for nuclear speckles, ribosomal, and other proteins. (Wilcoxon rank-sum test. ns, not significant.)

### RRP1B shows differential association with RNA transcripts after thermal stress

Based on the associations of RRP1B with rRNA processing and splicing from the RRP1B interactome (Fig. 3), RNA immunoprecipitation sequencing (RIP-seq) of nuclear RNA was performed to investigate potential changes in RRP1B associations under cellular stress. Since RRP1B has previously been primarily associated with non-coding RNA (rRNA), alignments were performed using the Telomere-to-telomere version 2 (T2T v2) genome to try to capture a more comprehensive set of potential non-coding associations, including ribosomal repeat regions. Comparing RRP1B IP at standard (37 °C) and heat shock (42 °C) revealed 1,594 transcripts (1,280 individual genes) displaying increased association with RRP1B and 4,838 transcripts (4,414 individual genes) with reduced association after thermal shock (Fig. 4A and Table S3). The majority of transcripts associated with RRP1B at both temperatures were protein-coding (Fig. S4A-B). Non-coding transcripts were overrepresented in the depleted transcripts (N = 515, 11.6%), compared to the enriched transcripts (N = 15, 1.1%) (Fig. 4B), suggesting RRP1B preferentially binds protein-coding transcripts after cellular stress. Unexpectedly, a significant difference in average gene size was also observed, with a median length of the enriched genes of 25,147 bp compared to 65,532 bp for the depleted class (p = 3.3E-133) (Fig. 4C). Analysis of the primary transcripts (Annotation using MANE Select, APPRIS principal_1) revealed that the canonical fully spliced mRNA median length of the depleted transcripts was 1.48-fold longer (3.9kb vs 2.6 kb, p=1.2E-47)(Fig. 4D) and that the median intron length of depleted genes was 2.34-fold longer than that of the enriched genes (3.5kb vs 1.5 kb, p=2.3E-89) (Fig. 4E). Recent studies have suggested a radial gradient of transcription within the nucleus, with shorter GC-rich genes and transcripts enriched in the nuclear interior near NS, where they are also associated with RNA splicing ontologies, whereas longer genes and transcripts are enriched at the nuclear periphery^4,6,7,41^. Consistent with this, the shorter RRP1B enriched transcripts were highly enriched in splicing ontologies (GO: p.adj < 2.2E-74) (Fig. 4F-G). Both enriched and depleted transcripts were enriched in transcription ontologies (GO: p.adj < 3.4E-74), concordant with the known effects of heat shock on cellular transcription (Fig. 4G and Fig. S4C). The RIP-seq data were therefore intersected with published studies examining subnuclear RNA localization using APEX-seq^7^, an RNA proximity labeling technique for spatial mapping of RNA organization. Comparison with APEX2-seq datasets targeting NS revealed that 25.39% of genes with RRP1B-enriched transcripts overlapped with NS-proximal APEX-seq transcripts, compared with 18.23% of genes with RRP1B-depleted transcripts (Fig. S4D and Table S4). Similar proportions of enriched and depleted transcripts overlapped with transcripts associated with the nuclear membrane (LMNA, ∼44%) and nucleoli (FBL, ∼30%) (Fig. S4E-F).

**Figure 4.**
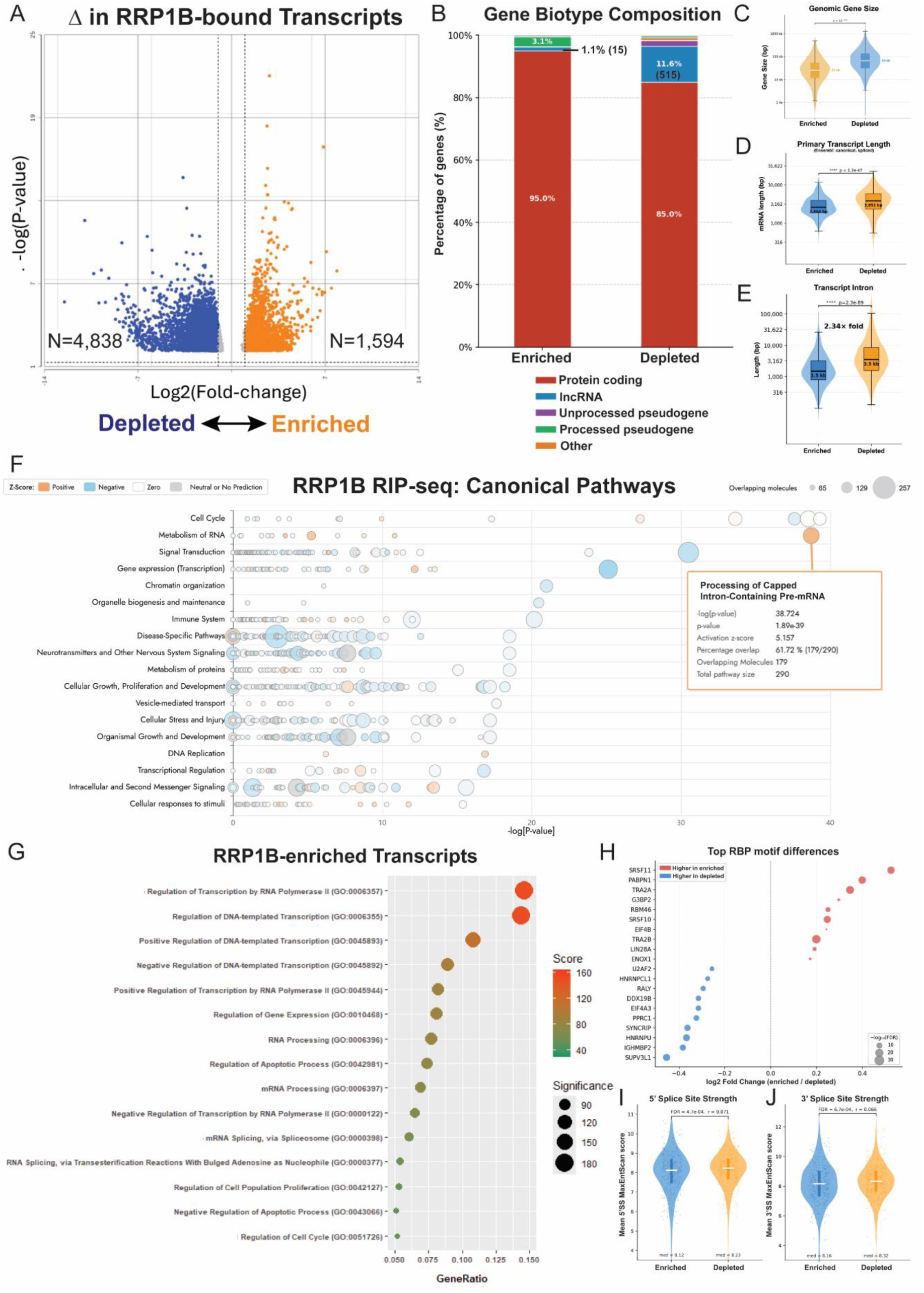
Differential RNA transcript association of RRP1B in response to thermal stress. (A) Volcano plot showing RRP1B-bound transcripts with significant changes upon stress. (B) Stacked percentage bar charts depicting the composition and relative abundance of RNA biotypes within RRP1B-enriched and RRP1B-depleted transcript populations identified through RIP-seq analysis. (C-E) Violin plots comparing gene length (C), primary transcript length (D), and transcript intron length (E) between RRP1B-enriched and RRP1B-depleted transcript populations. (Wilcoxon-Mann-Whitney test.) (F) Bubble chart overview of the top canonical pathways identified by Ingenuity Pathway Analysis (IPA) for RRP1B-associated transcripts in response to stress. (G) Dot plot of the top enriched Gene Ontology pathways among RRP1B-enriched transcripts under stress conditions. (H) Dot plot showing the top enriched RNA-binding protein (RBP) motifs identified from RIP-seq analysis. (I-J) Violin plots of 5′(I) and 3′(J) splice-site strength in RRP1B-enriched and RRP1B-depleted transcripts. (Two-sided Mann–Whitney U test with Benjamini–Hochberg FDR correction.)

RNA binding motif (RBP) analysis^42^ was then performed to identify any known binding motifs presenting in the RRP1B-enriched or depleted gene sets. Functional annotation of the RBPs with significantly higher motif density in RRP1B-depleted transcripts revealed enrichment for factors of late-stage nuclear RNA processing, including in co-transcriptional 3′ splice site recognition (U2AF2)^43^, exon junction complex deposition and NMD surveillance (EIF4A3)^44^, and mRNA export at the nuclear pore (DDX19B)^45^ (Fig. 4H). In contrast, RBPs enriched in RRP1B-associated transcripts (SRSF11, TRA2A, TRA2B, and SRSF10) are canonical SR proteins that bind exonic splicing enhancers and drive constitutive exon definition during early co-transcriptional splicing^46,47^ (Fig. 4H). Analysis of the splice sites present in the RIP-seq transcripts revealed that both the 5’ and 3’ splice site scores were significantly lower in the RRP1B-enriched transcripts compared to the depleted transcripts (p = 4.7E-4 and 6.7E-4, respectively), indicating that the enriched transcripts had, on average, slightly weaker canonical splice sites compared to the depleted genes (Fig. 4I-J).

Together, these data suggest that under standard tissue culture conditions, RRP1B is associated with longer genes positioned closer to the periphery of the nucleus^7,41^, potentially including at the RRP1B-SUN2 structures (Fig. 2B-F). Under thermal stress, however, RRP1B preferential associates with shorter, splicing-enriched transcripts positioned at or near the nuclear speckle, suggesting that RRP1B differentially associates with positionally distinct classes of RNA under cellular stress.

### RRP1B-associated transcripts are enriched for retained introns or skipped exons

The nuclear speckle is a membrane-less condensate associated with high concentrations of splicing factors, which has more recently been associated with both co-transcriptional and post-transcriptional splicing^6,48^. Due to the enrichment of RRP1B in the nuclear speckle after heat shock, differential splicing analysis of the RRP1B RIP-seq data was performed using the replicate multivariate analysis of transcript splicing (rMATS) algorithm. Due to incompatibility of the T2Tv2 annotations, rMATS analysis was performed using the hg38 genome. The most frequent alternative splicing events observed were skipped exons (SE) or retained introns (RI) (Table S5). Ontology analysis demonstrated that SE splicing changes were primarily involved in the regulation of protein/macromolecule metabolic processes and post-translational modification (Fig. 5A), whereas transcripts with RI events were enriched in gene expression and RNA splicing and RNA nuclear export ontologies (Fig. 5B). Consistent with these findings, transcripts of several canonical SR proteins and RNA export factors identified among the RI events exhibited reduced RRP1B binding to intronic regions under stress conditions (Fig. 5C and Fig. S5A). This suggested that RRP1B may contribute to the previously described retained intron autoregulatory feedback mechanisms of splicing and RNA export factors^49–51^.

**Figure 5.**
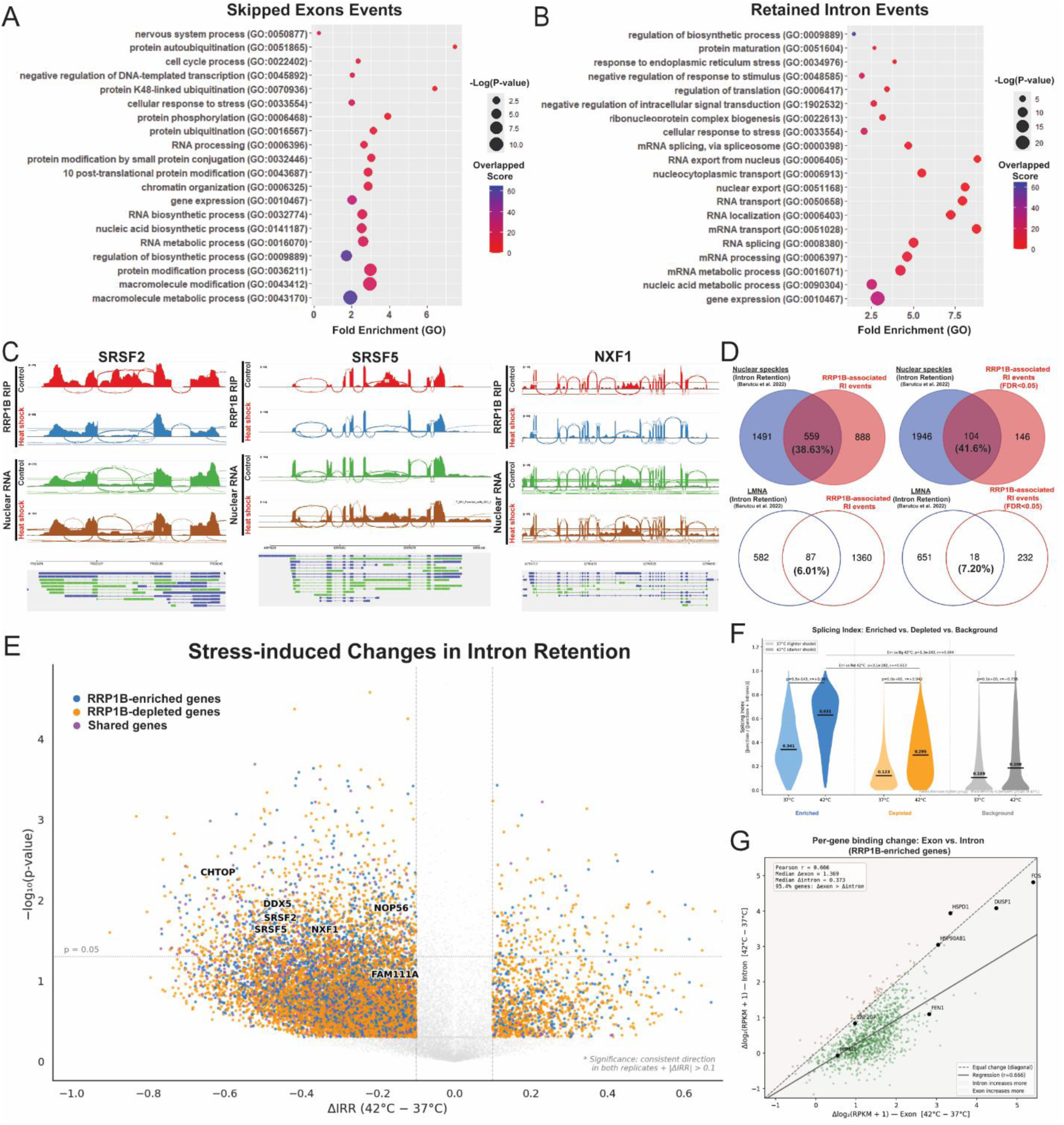
Alternative splicing events, including retained introns and exon skipping, are enriched in RRP1B-associated transcripts. (A-B) Dot plots of top Gene Ontology enrichments for RRP1B-associated exon skipping (A) and intron retention (B) splicing events. (C) Sashimi plots illustrating several representative transcripts involved in significant RRP1B-associated retained-intron (RI) splicing events. (SRSF: Serine/arginine-rich splicing factor; NXF1: Nuclear RNA export factor 1) (D) Venn diagrams showing overlapping RI events among NS APEX-seq, nuclear membrane (LMNA) APEX-seq, and stress-induced RRP1B-associated RI events. (E) Volcano plot of stress-induced differential RRP1B binding at RI. (ΔIRR: Intron retention ratio) (All label genes are involved in significant RRP1B-associated RI splicing events) (F) Violin plot of splicing indices for RRP1B-enriched, RRP1B-depleted, and unbound transcripts at 37 °C and 42 °C. (Paired Wilcoxon signed-rank test.) (G) Scatter plot of RRP1B binding changes in exon and intron regions between 37 °C and 42 °C.

Since the nuclear speckle has previously been associated with retained introns, we reanalyzed the APEX-seq^7^ data specifically for RI transcripts that revealed 20.0% of genes with RRP1B-enriched transcripts and 10.51% RRP1B-reduced transcripts under heat shock overlapped with retained-intron transcripts proximal to NS. (Fig. S5B-C and Table S4). In contrast, only ∼4% of genes in RRP1B-associated transcripts overlapped with transcripts associated with nuclear membranes (LMNA) (Fig. S5B-C). Moreover, 38.63% RRP1B-associated RI splicing genes identified by rMATS analysis (Table S4) overlapped with NS APEX2-seq RI transcripts, whereas only 6.01% overlapped with LMNA APEX2-seq RI transcripts (Fig. 5D). Comparison with another public dataset^4^ using the reverse transcribe and tagment (RT&Tag) approach to detect RNAs in NS also showed a 45.54 % overlap between RRP1B-associated RI splicing genes and NS-localized RI transcripts^4^ (Fig. S5D-E and Table S4). Analysis of RRP1B-bound introns using the intron retention rate (IRR), defined as the fraction of intronic reads among total exonic and intronic reads, revealed that heat shock dramatically reduced intron retention in RRP1B-associated RNA, with 10,180 introns showing decreased retention and 1,337 showing increased retention (Fig. 5E). Interestingly, RRP1B-enriched transcripts exhibited a significant increase in splicing index under stress conditions and displayed significantly higher splicing indices than both RRP1B-depleted transcripts and background transcripts that were not significantly bound by RRP1B (Fig. 5F). Further analysis of RRP1B-bound exonic and intronic regions within the enriched transcripts revealed a preferential increase in RRP1B binding to exons under heat shock conditions (42 °C) relative to control conditions (37 °C) (Fig. 5G and Fig. S5F-G).

Collectively, these findings suggest that RRP1B may redistribute from nuclear membrane–associated complexes to NS–associated splicing and transcriptional hubs under cellular stress, where it may participate in regulating RI splicing and consequently exhibit preferential association with exon-dense, intron-poor transcripts.

### RRP1B overexpression reduces retained-intron transcripts in the cytoplasm

To better understand the role of RRP1B in RNA processing and the fate of transcripts differentially bound by RRP1B under stress conditions, RNA-seq analysis was performed on cytoplasmic and nuclear fractions of MDA-MB-231 cells, with or without RRP1B overexpression (oeRRP1B), under either 37 °C or 42 °C conditions (Fig. S6A–C). The effect of oeRRP1B on splicing events was then evaluated across each condition and subcellular fraction relative to parental cells. RRP1B overexpression induced significant alterations in alternative splicing across all four fractionation conditions, with the magnitude and directionality of effects varying by compartment and temperature. In the cytoplasm at 37°C, the most striking finding is a nearly total suppression of RI, with 89.4% of significant RI events show decreased retention in oeRRP1B cells (binomial p < 1E-117) (Fig. 6A and Fig. S6D), suggesting RRP1B promotes either the splicing or nuclear retention of RI-containing transcripts under basal conditions. Skipped exon events show the opposite trend in the cytoplasm, with oeRRP1B favoring increased exon skipping, which increases under heat stress (Fig. 6A and Fig. S6D). In the nucleus at baseline (37°C), oeRRP1B has minimal directional impact on most event types, with only SE events showing a significant bias, indicating that the impact of RRP1B on splicing at steady state is primarily manifest in the cytoplasmic fraction. Under heat stress, however, in the nuclear compartment oeRRP1B shifts the balance toward decreased inclusion across all types of splicing events (22–42% in OE) (Fig. 6A and Fig. S6D), in accordance with enhanced nuclear splicing or export of these transcripts during the stress response^52,53^. Collectively, these data indicate that RRP1B shapes the splicing landscape in a compartment- and stress-dependent manner, with its most significant effects on retained introns in the cytoplasm.

**Figure 6.**
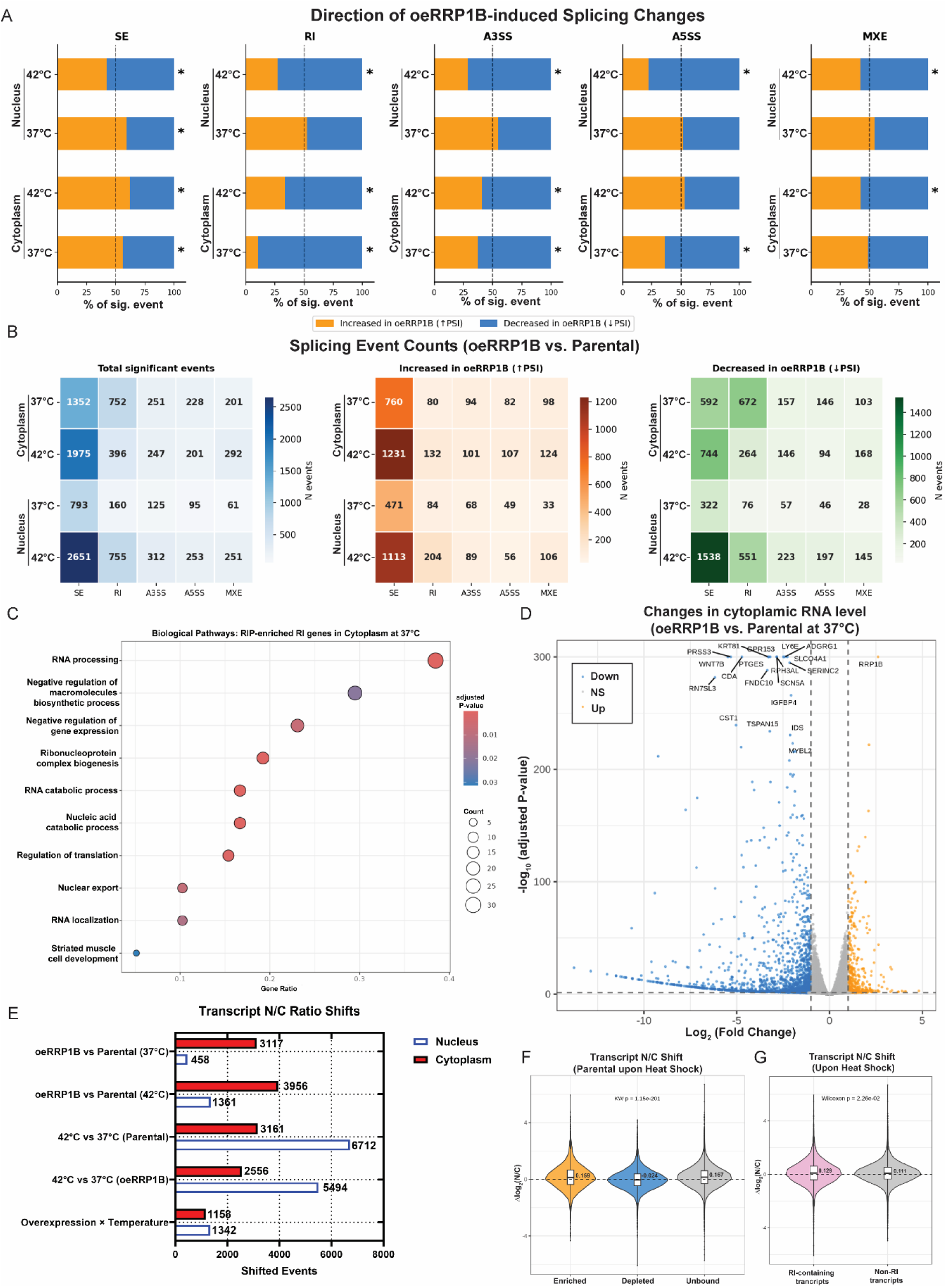
RRP1B overexpression decreases cytoplasmic retained-intron transcripts. (A) Stacked percentage bar charts showing the distribution of differential splicing events, including skipped exons (SE), retained introns (RI), alternative 3′ splice sites (A3SS), alternative 5′ splice sites (A5SS), and mutually exclusive exons (MXE), across different subcellular fractions and temperature conditions following RRP1B overexpression. (*, p < 0.05 significant directional bias, binomial test.) (B) Heatmaps of directional splicing changes induced by RRP1B overexpression across subcellular fractions and temperature conditions. (C) Dot plot of top Gene Ontology enrichments for cytoplasmic RRP1B-enriched RI genes under normal culture conditions. (D) Volcano plot illustrating differential cytoplasmic RNA expression induced by RRP1B overexpression. (E) Bar chart showing transcript nuclear-to-cytoplasmic (N/C) ratio shifts across multiple comparison conditions. (F) Violin plot of transcript N/C ratio shifts in RRP1B-enriched, RRP1B-depleted, and unbound transcripts in parental cells. (Kruskal–Wallis test.) (G) Violin plot of transcript N/C ratio shifts in RI-containing, non-RI transcripts in parental cells. (Wilcoxon test.)

### RRP1B binding correlates with differential splicing of specific event types

To determine whether RRP1B-bound transcripts are preferentially subject to oeRRP1B-induced splicing changes, significant splicing events were filtered for RRP1B RIPseq-enriched and - depleted gene lists. RIP-enriched genes (those gaining RRP1B binding under heat stress; N = 1,280) and RIP-depleted genes (those losing binding under heat stress; N = 4,414) were each tested for over-representation among differentially spliced genes using Fisher’s exact tests with Benjamini-Hochberg FDR correction across all 40 contrast-event type combinations (Fig. 6B). RIP-depleted genes were significantly over-represented among oeRRP1B-induced skipped exon changes in all four compartment/temperature conditions (Fisher’s OR 1.2–1.6, FDR = 2E-6 to 9E-4), indicating that RIP-depleted transcripts are most sensitive to RRP1B level–dependent exon skipping. Ontology analysis of the 484 genes with SE events in the 42 °C cytoplasmic parental versus oeRRP1B revealed no significantly enriched ontology after multiple testing correction (GO: BP, FDR-adjusted p = 9.7E-1). In contrast, RIP-enriched genes were over-represented among cytoplasmic retained intron events at baseline temperature (OR = 1.73, FDR = 8.8E-4) and were highly enriched for genes associated with RNA processing (GO: BP, FDR-adjusted p = 8.03E-9) (Fig. 6C). No significant splicing was detected for A3SS, A5SS, or MXE events for RRP1B RIP-enriched or depleted genes, after FDR correction, indicating these event types are regulated by oeRRP1B through indirect or RRP1B-independent mechanisms. These suggest that transcripts gaining RRP1B binding under heat stress are subject to RI regulation and are consistent with the role of RRP1B in the regulation of cellular splicing in response to microenvironmental stress.

### RRP1B regulates nuclear retention of its associated transcripts during heat shock

Transcript-level quantification of nuclear and cytoplasmic RNA-seq fractionation data from parental and RRP1B-overexpressing MDA-MB-231 cells was performed to investigate how RRP1B affected processing and nuclear export. Results showed overexpression of RRP1B induced substantial transcriptional reprogramming (Fig. 6D and Fig. S6E-G). In the cytoplasmic compartment at 37°C, 1,666 genes were differentially expressed (422 up, 1,244 down; adjusted p < 0.05, |log2FC| ≥ 1) with 1,438 cytoplasmic DEGs were identified (347 up, 1,091 down) at 42°C (Fig. 6D). In the nuclear fractions, 835 DEGs were detected at 37°C (121 up, 714 down) and 1,143 at 42°C (297 up, 846 down) (Fig. S6E-F). Across all four comparisons, downregulated genes consistently outnumbered upregulated genes by 3- to 6-fold, consistent with a role for RRP1B in promoting RNA stability or nuclear export. Nuclear/Cytoplasmic (N/C) ratio analysis revealed that RRP1B overexpression preferentially drives cytoplasmic accumulation of a certain subset of transcript isoforms (Fig. 6E). At 37°C, 3,117 transcripts shifted toward the cytoplasm upon RRP1B overexpression versus only 458 that shifted toward the nucleus (p < 0.05, |Δlog2(N/C)| ≥ 0.5). This cytoplasmic bias was amplified at 42°C, with 3,956 cytoplasmic and 1,361 nuclear shifts. As expected, heat shock itself induced a pronounced nuclear retention response: in parental cells, 6,712 transcripts shifted toward the nucleus versus 3,161 toward the cytoplasm upon heat shock, consistent with stress-induced nuclear retention of incompletely processed transcripts. A similar pattern was observed in oeRRP1B cells (5,494 nuclear vs. 2,556 cytoplasmic shifts). The interaction analysis identified 4,068 transcripts with temperature-dependent changes in RRP1B-driven localization (nuclear shift) and 2,041 with cytoplasmic shifts, indicating that a substantial effect of RRP1B on subcellular RNA localization is modulated by thermal stress.

To investigate whether RRP1B binding status affects transcript localization dynamics, Kruskal-Wallis tests across three binding categories (enriched, depleted, unbound) were performed. Shifts between nucleus and cytoplasmic compartments were highly significant for all five N/C comparisons (oeRRP1B vs Parental, 37°C, oeRRP1B vs Parental, 42°C, 42°C vs 37°C in Parental cells, 42°C vs 37°C in oeRRP1B cells, overexpression status × temperature interaction, p = 2.1E-8 to 1E-300) (Fig. 6F and Fig. S6H). Under heat shock, RRP1B-enriched transcripts shifted toward the nucleus (mean Δlog2(N/C) = +0.159 in parental cells, +0.178 in oeRRP1B cells), while RRP1B-depleted transcripts showed a modest but significant shift toward the cytoplasm (mean Δlog2(N/C) = −0.024 and −0.029, respectively) (Fig. 6F and Fig. S6H). This suggests that RRP1B binding during heat shock promotes nuclear retention of its RNA targets, while transcripts that lose RRP1B association are preferentially exported. Notably, RRP1B-bound genes were significantly underrepresented among steady-state DEGs across all four compartment/temperature comparisons (Fisher’s exact test, OR = 0.23–0.50, p < 1E-10), consistent with RRP1B’s primary regulatory impact is on RNA localization rather than transcriptional output or RNA stability (Fig. S6I). Integration with rMATS-derived retained intron data (49,659 RI events across eight comparisons; 1,943 RI-containing genes at FDR < 0.05) also revealed significant N/C ratio shifts between RI-containing and non-RI transcripts across all five comparisons (Wilcoxon rank-sum test, p = 9.3E-7 to 5.8E-17). Under RRP1B overexpression at 37°C, transcripts from RI-containing genes showed a mean Δlog2(N/C) of −0.188 compared to −0.204 for non-RI transcripts (p = 9.3E-7) (Fig. S6J), indicating that RI-containing transcripts are less cytoplasmic under conditions of higher RRP1B expression. Under heat shock, RI-containing transcripts shifted toward the nucleus to a similar degree as non-RI transcripts (mean Δlog2(N/C) +0.129 vs. +0.111, p = 0.023) (Fig. 6G), suggesting that the nuclear retention response by heat shock is broadly distributed across the transcriptome regardless of splicing status.

### RRP1B selectively facilitates the nuclear export of protein-coding mRNA under cellular stress conditions

Given our findings that differential RRP1B binding to transcripts with or without retained introns correlates with transcript subcellular distribution under stress conditions, transcript levels of protein-coding and Ensembl-annotated RI isoforms from all genes, as well as the 250 RI genes identified by rMATS analysis, were further compared across all compartments and temperature conditions. The significant increase in cytoplasmic export of fully spliced protein-coding isoforms among the 250 RI genes, a trend also observed across all genes, together with the absence of a corresponding increase in nuclear retention of retained-intron isoforms (Fig. 7A-B), an effect further enhanced by RRP1B overexpression (Fig. 7C and Fig. S7A), suggests that RRP1B facilitates the export of these transcripts following its association with their retained-intron counterparts under stress conditions. Under normal culture conditions, distribution of nuclear and cytoplasmic RNA for protein-coding isoforms was not significantly different from that of RI transcripts (Fig. 7D and Fig. S7B). Under heat shock conditions, more protein-coding transcripts, especially those associated with RRP1B (ΔFold = 2.85 for all genes, ΔFold = 3.22 for RRP1B-associated RI transcripts, Fig. 7E-F), were enriched in the cytoplasmic fraction, suggesting that RRP1B selectively promotes the nuclear export of protein-coding transcripts under stress. These findings are consistent with our current RRP1B IP-MS data (Fig. 3 and Table S2) and previous studies^36^ identifying Aly/REF Export Factor (ALYREF/THOC4), a nuclear mRNA export adaptor, as an RRP1B-interacting protein, supporting a functional association between RRP1B and transcript transport. One striking example was SRSF2, a well-known RNA splicing factor. SRSF2-202, the major protein-coding isoform, exhibited significantly reduced nuclear levels and increased cytoplasmic abundance upon heat shock in both parental and RRP1B-overexpressing MDA-MB-231 cells (Fig. 7G). In contrast, the RI-containing isoform, SRSF2-209, showed increased nuclear retention under heat shock conditions (Fig. 7G). A similar pattern was observed for NXF1, an RNA export factor, in which the major protein-coding transcript NXF1-201 displayed increased cytoplasmic abundance, whereas RI-containing isoforms, including NXF1-205 and NXF1-215, remained minimally expressed in the cytoplasm (Fig. S7C). Overall, lower cytoplasmic levels of RRP1B-associated RI transcripts were detected, accompanied by increased abundance of the corresponding fully spliced transcripts under stress conditions (Fig. 7H). Collectively, these findings demonstrate that RRP1B promotes retention of RI transcripts in the nucleus and selectively export of protein-coding isoforms, suggesting RRP1B plays a role in regulating intron excision under stress conditions.

**Figure 7.**
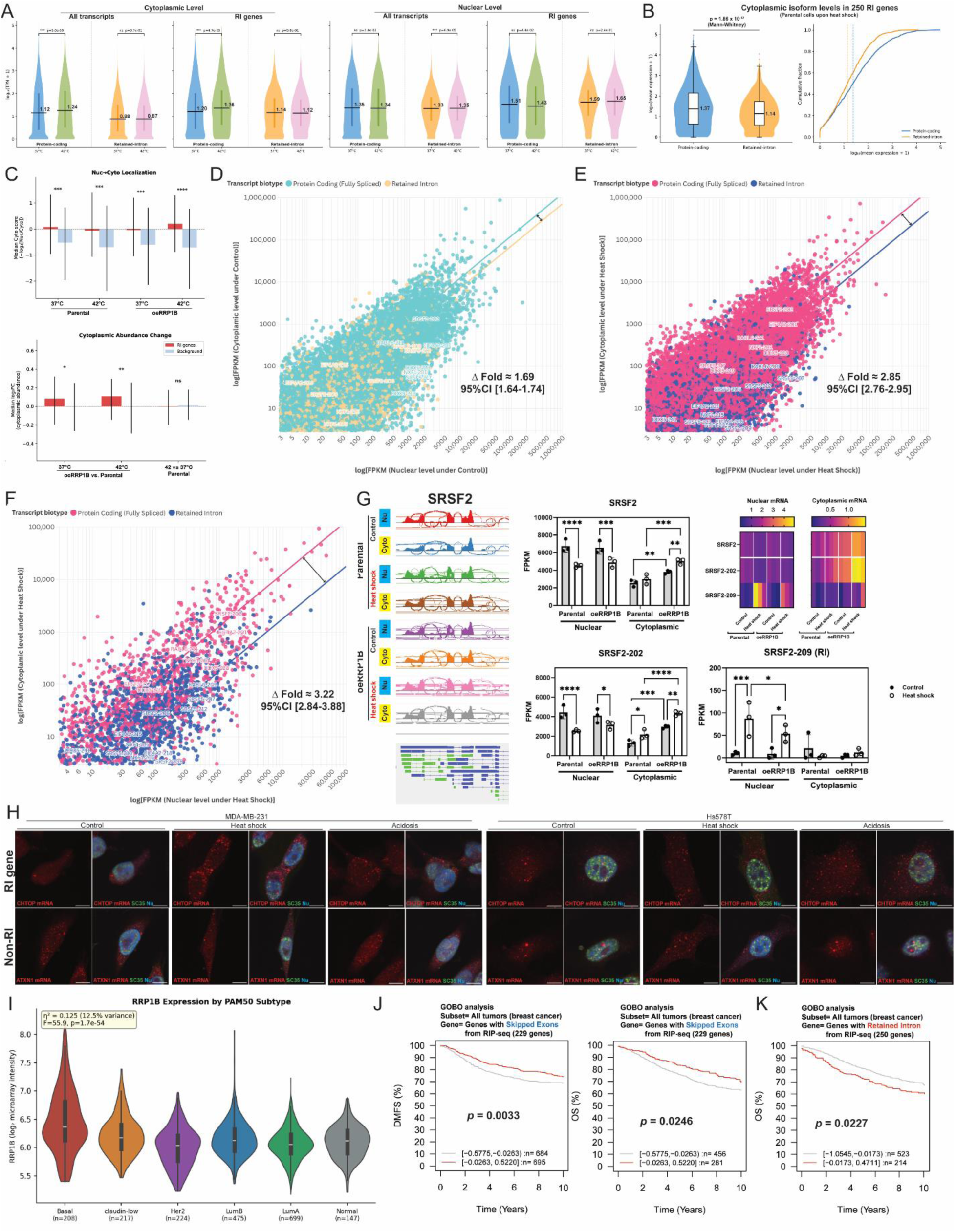
Selective export of protein-coding mRNAs by RRP1B under cellular stress conditions. (A) Violin plots comparing protein-coding and retained-intron (RI) isoforms across subcellular fractions for all genes and the 250 RI genes. (Mann-Whitney U test.) (B) Violin (left) and ECDF (right) plots comparing cytoplasmic RNA levels between all genes and the 250 RI genes in parental cells under heat shock. (Mann–Whitney test for violin plots; Wilcoxon signed-rank test for ECDF plots.) (C) Bar plots of Nuc/Cyto expression ratios and localization shifts for the 250 RI genes. (Mann-Whitney U test. *, q<0.05, **, q<0.001, ***, q<1E-6, ****, q<1E-10) (D) Scatter plot with trend lines showing the ratio between nuclear and cytoplasmic levels in protein-coding and RI transcripts in MDA-MB-231 parental cells under normal culture conditions. (CI: Confidence Interval) (E) Scatter plot with trend lines showing the ratio between nuclear and cytoplasmic levels in protein-coding and RI transcripts in MDA-MB-231 parental cells under heat shock conditions. (F) Scatter plot with trend lines showing the ratio between nuclear and cytoplasmic levels in protein-coding and RI transcripts among 250 RI genes in MDA-MB-231 parental cells under heat shock conditions. (G) Sashimiplots, quantitative analyses, and heatmaps showing the changes in RRP1B-binding and subcellular RNA expression of SRSF2 transcript isoforms upon stress. (Two-way ANOVA with Fisher’s LSD test. *, p<0.05, **, p<0.01,***, p<0.0005, ****, p<0.0001) (H) Representative RNA-FISH images of RI and non-RI genes in MDA-MB-231 and Hs578T cells treated with heat shock (42°C) for 3 hours, acidic shock (pH6.5) for 1 hour, or under normal culture conditions (37°C, pH7.4). Scale bar, 10 μm. (I) Violin plots of the RRP1B-associated transcriptomic signature across METABRIC PAM50 subtypes. (J) GOBO (Gene expression-based Outcome for Breast cancer Online; LUND University) analysis of 229 genes of RRP1B-associated SE transcripts in Distant Metastasis-Free Survival (DMFS) and overall survival (OS) in breast cancer patients. (K) GOBO analysis of 164 genes of RRP1B-associated RI transcripts in overall survival (OS) in breast cancer patients.

### RRP1B promotes a basal-like transcriptomic program, and its stress-associated alternative splicing signature strongly impacts patient outcomes

To assess whether the transcriptional program induced by RRP1B overexpression in MDA-MB-231 cells is reflected in primary breast tumors, we generated a weighted signature score from the top 25 upregulated and top 25 downregulated cytoplasmic DEGs at 37 °C and applied it to 1,974 METABRIC breast tumors. Of the 50 signature genes, 37 were represented on the microarray platform. The signature varied significantly across PAM50 subtypes (ANOVA p = 7.2E-149), with Basal-like tumors showing the highest scores, followed by claudin-low tumors, with luminal, HER2-enriched, and normal-like tumors exhibited substantially lower scores (Figure. 7I). The signature distinguished Basal from non-Basal tumors with an AUC of 0.849 and TNBC from non-TNBC tumors with an AUC of 0.828, while HER2-enriched tumors were poorly separated from luminal subtypes (AUC = 0.599) (Fig. S7D). The strongest contributors to subtype separation were SPDEF, AFF3, and AGR2, luminal-associated genes strongly downregulated by RRP1B overexpression and correspondingly reduced in Basal tumors within the METABRIC cohort.

To further determine the clinical significance of RRP1B-mediated splicing regulation, Kaplan–Meier survival analysis was performed using the RRP1B-associated genes from the RIP-seq data that showed significant differences in SE (229 genes) or RI (250 genes) splicing events as independent gene signatures (Table S5 and Fig. 5A-B). Gene expression-based Outcome for Breast cancer Online (Gobo) analysis showed high expression of the SE signature led to an improved prognosis in breast cancer (Fig. 7J). On the other hand, high level of the 250 RI gene set was significantly associated with worse overall survival, with stronger impact shown in higher grades of breast cancers (Fig. 7K and Fig. S7E). This suggests that RRP1B plays a critical role in regulating the splicing of a subset of genes that are essential for maintaining cancer cell survival under stress conditions.

In summary, in addition to its previously described role in ribosomal biogenesis, these data suggest that RRP1B plays an important role in cancer cellular response to microenvironmental stresses. Through a combination of increased protein production and/or stability and changes in subnuclear translocation, RRP1B helps regulate the splicing and nuclear retention/export of distinct classes of RNAs, particularly those associated with retained introns and skipped exons. Increased expression of RRP1B results in export into the cytoplasm of fully spliced transcripts associated with worse patient outcome and RNA processing and splicing and induces a more basal-like transcriptional pattern. Taken together, the data highlight a novel, microenvironmentally sensitive stress mechanism associated for RRP1B in the malignant progression of breast cancer.

## Discussion

Previously, our laboratory identified RRP1B as an inherited metastasis susceptibility gene for breast cancer^54^. However, the mechanism by which RRP1B acted in the metastatic process has been unclear. Immunofluorescence analysis has revealed that under standard tissue culture conditions RRP1B is primarily localized within the nucleolus (Ex. Human Protein Atlas https://www.proteinatlas.org/). RRP1B contains an N-terminal NOP52 domain which has previously been associated with pre-rRNA cleavage and processing^55^. Consistent with this, RRP1B has previously shown to play an important role in pre-rRNA processing and the nuclear export of the large ribosomal subunit^56^. However, results from related studies in our laboratory have not revealed a significant role for ribosomal biogenesis in our experimental metastasis assays^57^. Proteomics studies, both previous and current, have also revealed an association between RRP1B both splicing factors^58,59^ and nuclear export factors^54^, suggesting additional roles for RRP1B in transcript processing. Work from other laboratories have demonstrated that RRP1B contributes to alternative splicing^58^, though the mechanism through this might contribute to tumor progression was not clearly defined.

Here we demonstrate that RRP1B functions within tumor cells as part of a microenvironmental stress response pathway. Under tumor-relevant stresses, including increased extracellular stiffness and temperature^60^, hypoxia^61^, acidosis^62^, RRP1B disassociates from transcripts of longer genes that are preferentially positioned towards the nuclear periphery. Under these conditions, RRP1B concentrates in the nuclear speckle and becomes associated with shorter transcripts enriched for splicing and RNA processing factors. This redistribution alters the nuclear-to-cytoplasmic balance of RRP1B-associated transcripts, with transcripts depleted from RRP1B showing modest cytoplasmic accumulation, whereas transcripts gaining association with RRP1B are preferentially retained in the nucleus during stress. Importantly, fully spliced isoforms derived from the subset of transcripts undergoing RRP1B-associated retained-intron processing exhibit increased cytoplasmic abundance under stress conditions. These findings suggest that RRP1B plays a critical role in rapidly remodeling the transcriptome during the early stage of metastatic tumor growth and progression, enabling cancer cells to survive and disseminate from the primary tumor as intratumoral conditions become increasingly inhospitable.

The molecular basis of stress-induced NS localization maps to a set of phosphorylation events within the IDRs of the C-terminal domain of RRP1B. The finding that phosphomimetic substitutions at S245, S350, S702, and S706 promote NS-proximal puncta formation or large peripheral aggregates, while unphosphorylatable alanine substitutions confine the protein to the nucleolus, is consistent with a model in which IDR phosphorylation tunes the phase-separation propensity of RRP1B and its partitioning between condensate compartments^27,63^. The sensitivity of puncta to RNase treatment further suggests that RNA scaffolding contributes to the assembly of these structures, consistent with emerging models of RNA-seeded condensate nucleation. The distinct behaviors of S702A versus S702D suggested that the phosphomimetic disrupts both nucleolar PP1β/γ interaction and SUN2-associated nuclear envelope localization, and that stress-induced phosphorylation releases it for redistribution to the NS. The partial phenotypes of S245D and S350D — with cell-to-cell variability between wild-type and aggregated patterns — suggest that no single phosphorylation event is sufficient and that combinatorial modification is required for full NS integration, consistent with the multi-site phosphorylation logic increasingly recognized for IDR-containing condensate components.

In addition to the role in modulating RNA nuclear export, RRP1B contributes to cellular stress response through modulation of retained introns. Originally thought to be the result of inefficient splicing, retained introns are now thought to play an important role in a variety of biological processes, including stress responses^64,65^. Within the nucleus, RRP1B disassociates with retained intron-containing transcripts, for example, SRSF splicing family transcripts, and becomes exclusively associated with fully spliced messages. The majority of RI-containing transcripts are also lost in the cytoplasmic fraction, suggesting that RRP1B may be retaining transcripts within the nucleus to permit the completion of less-efficient splicing before export to the cytoplasm. As mentioned above, these transcripts are enriched in RNA processing and splicing factors, indicating that RRP1B is contributing to the previously described positive feedback loop modulating splicing factor production under stress^66^. In addition, the transcripts that lose RI in the cytoplasm with RRP1B overexpression predict poor outcome in breast cancer, consistent with higher expression of RRP1B itself being a poor prognostic marker and previous studies showing breast tumors with lower RI having worse outcome^67^. Taken together, these data indicate that RRP1B is likely contributing to tumor progression through retained intron-mediated adaptation to stress at the primary tumor site at the early steps of the metastatic cascade.

## Limitations

Several questions remain to be addressed. At present, it is not clear whether the source of RRP1B that accumulates in the nuclear speckle is from the nuclear envelope pools, the nucleolus, or from another as yet to be defined nuclear pool. Although the current study detected reduced RRP1B association with ribosomal proteins under stress conditions, suggesting that a subset of RRP1B-depleted transcripts may reside in the nucleolus, technical limitations prevented definitive confirmation of this observation. In particular, our inability to completely eliminate nucleolar localization of RRP1B, likely due to the inherently disordered nature of the protein, limited our ability to directly test this hypothesis. In addition, the molecular mechanism by which RRP1B localization is regulated requires further clarification. The data presented here suggest that serine phosphorylation, including the sites previously associated with PP1-mediated nucleolar stress response^56,68^, play an important role in RRP1B localization. However, due to the multiple phosphorylation sites present within the C-terminal IDR and aggregation and probable phase separation observed in the transient transfection assays, a complete understanding of the localization code remains to be elucidated. A comprehensive understanding of this mechanism likely will require more sophisticated methods, such as inducible systems or genome editing, to prevent inappropriate phase separation of phosphomimetic mutation constructs.

## Acknowledgements

The authors thank the Center for Cancer Research (CCR) Laboratory of Cancer Biology and Genetics (LCBG), CCR Confocal Microscopy Core Facility Head and staff members, Michael J. Kruhlak, Langston Lim, and Andy Tran, for technical assistance in microscopy analysis, CCR Genomics Core facility members Liz Conner and Madeline Wong, CCR sequencing facility (Frederick, MD) for RIP-seq/RNA-seq analysis. This work utilized the computational resources of the NIH HPC Biowulf cluster (https://hpc.nih.gov). The study was supported by the Intramural Research Program, National Cancer Institute, National Institutes of Health. The contributions of the NIH authors were made as part of their official duties as NIH federal employees, are in compliance with agency policy requirements, and are considered Works of the United States Government. However, the findings and conclusions presented in this paper are those of the authors and do not necessarily reflect the views of the NIH or the US Department of Health and Human Services.

## Author contributions

W.N.L. and K.W.H. conceived the project and designed experiments. W.N.L. performed the majority of the experiments and analyzed the data. K.W.H., M.P.L., and W.N.L. performed bioinformatics analysis. R.A. and M.J.K. provided research ideas and technical guidance. T.A. performed mass spectrometry analysis and analyzed the data. R.M.S. and J.S. performed immunohistochemistry staining, image analysis, and data interpretation. W.N.L. and K.W.H. wrote the manuscript.

---

**Figure S1.**
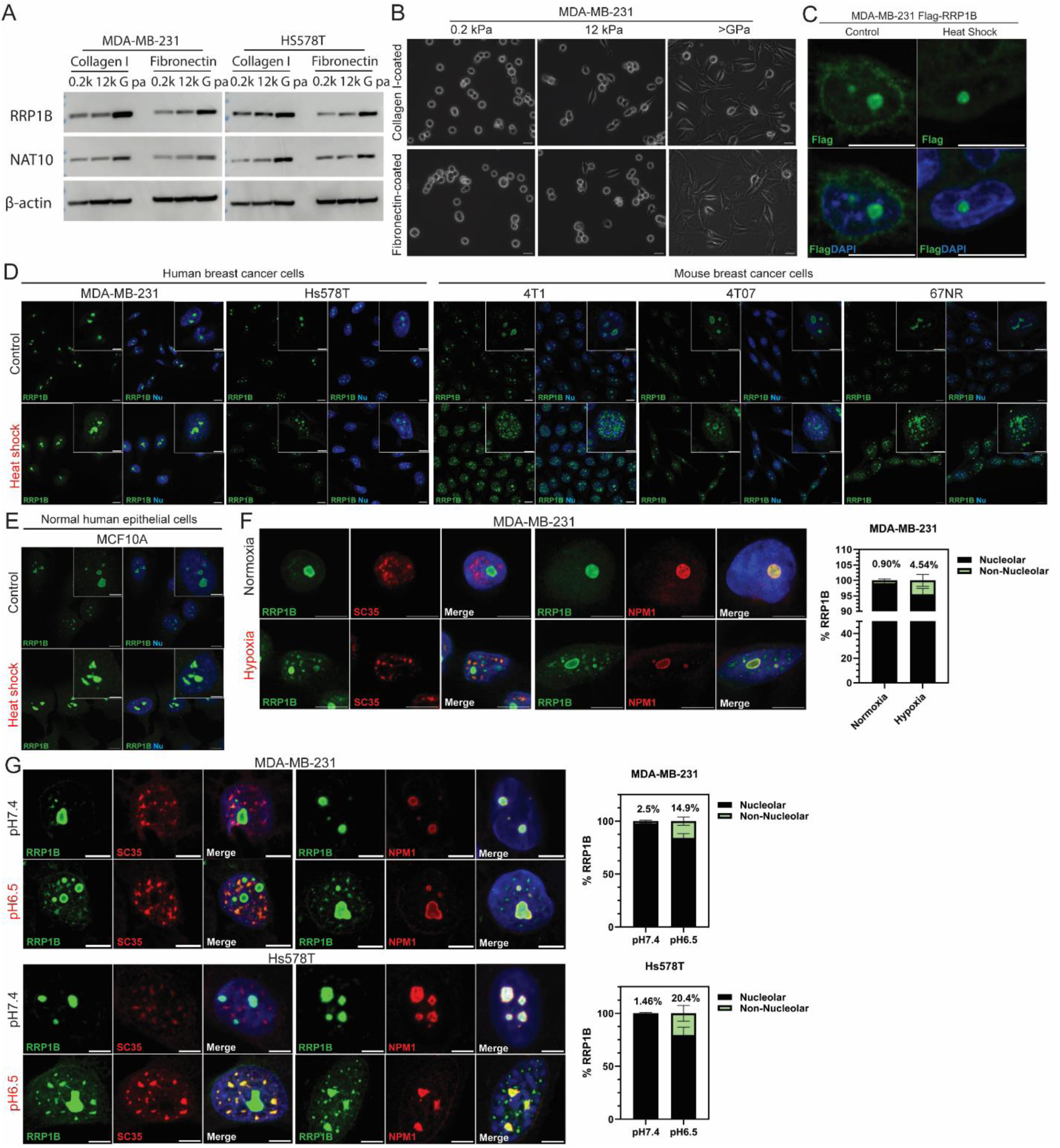
RRP1B expression and subnuclear localization under tumor-associated microenvironmental stresses, related to Figure 1. (A) Representative western blot images of both MDA-MB-231 and Hs578T cells seeded on hydrogels of different stiffness (0.2 kPa, 12 kPa, or >G Pa) that were coated with different extracellular matrix proteins (collagen I or fibronectin).(NAT10: N-Acetyltransferase 10, also a previously identified metastasis susceptibility complex partner.^69^)(B) Representative bright-field images of MDA-MB-231 cells seeded on hydrogels of different stiffness (0.2 kPa, 12 kPa, or >G Pa) that were coated with either collagen or fibronectin. Scale bar, 10 μm. (C) Representative immunofluorescence staining images of MDA-MB-231 cells stably expressing Flag-RRP1B cultured at either 37°C or 42°C for 3 hours. Scale bar, 10 μm. (D) Representative immunofluorescence staining images of human breast cancer cells (MDA-MB-231 and Hs578T), and mouse breast cancer cells (4T1, 4T07, and 67NR) cultured at either 37°C or 42°C for 3 hours. Scale bar, 5 μm. (E) Representative immunofluorescence staining images of normal human breast epithelial cells (MCF10A) cultured at either 37°C or 42°C for 3 hours. Scale bar, 5 μm. (F) Representative immunofluorescence staining images and quantitative analysis of MDA-MB-231 and Hs578T cells cultured in the medium with either 21% (Normoxia) or 1% (Hypoxia) oxygen for 1 hour. Scale bar, 5 μm. (G) Representative immunofluorescence staining images and quantitative analysis of MDA-MB-231 and Hs578T cells cultured in the medium with either pH7.4 or pH6.5 for 1 hour. Scale bar, 5 μm.

**Figure S2.**
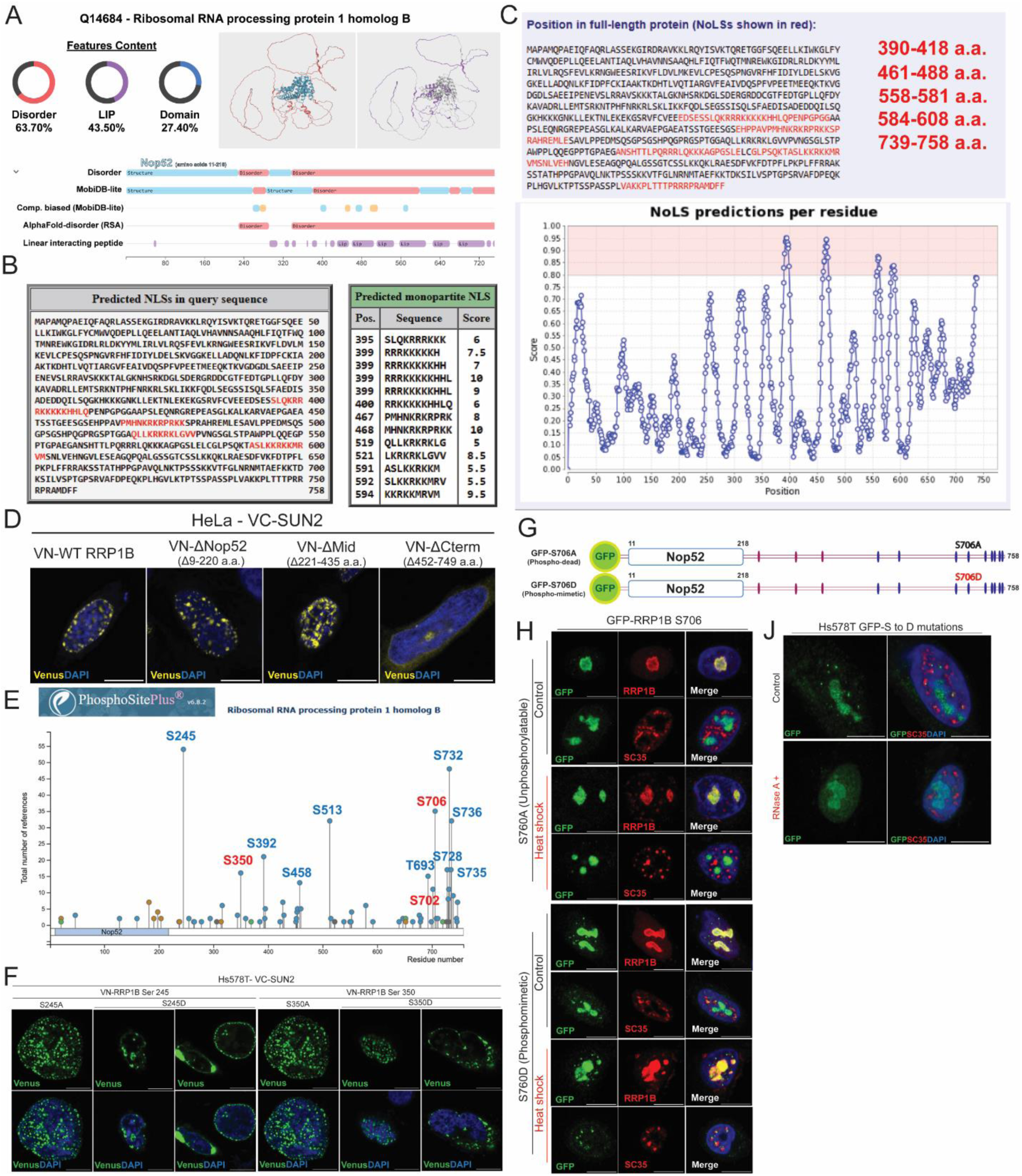
Characterization of RRP1B subnuclear localization dynamics through bioinformatic and mutation-based approaches, related to Figure 2. (A) MobiDB-based prediction of the RRP1B protein sequence showing intrinsically disordered regions (IDRs) and linear interacting peptides (LIPs). (B) Predicted nuclear localization signals (NLSs) in RRP1B identified using NLS Mapper. (C) Nucleolar localization signal (NoLS) prediction for RRP1B using the Nucleolar Localization Sequence Detector. (D) Representative BiFC images of HeLa cells co-transfected with VC-SUN2 and VN-RRP1B deletion constructs. Scale bar, 10 μm. (E) Annotated post-translational modification sites within RRP1B obtained from the PhosphoSitePlus database. (F) Representative BiFC images of HS578T cells co-transfected with VC-SUN2 and VN-RRP1B containing either the S245A or S350A (unphosphorylatable) or S245D or S350D (phosphomimetic) mutation. Scale bar, 10 μm. (G) Schematic representation of the RRP1B point mutation constructs used to determine the residues regulating subnuclear localization. (H) Representative immunofluorescence images of HS578T cells expressing GFP-tagged S706 A (unphosphorylatable) or S706D (phosphomimetic) RRP1B proteins cultured at either 37°C or 42°C for 3 hours. Scale bar, 10 μm. (J) Representative immunofluorescence images of HS578T cells expressing GFP-tagged D mutations treated with either vehicle or RNase A (0.1 μg/μL) at 37°C for 1 hour.

**Figure S3.**
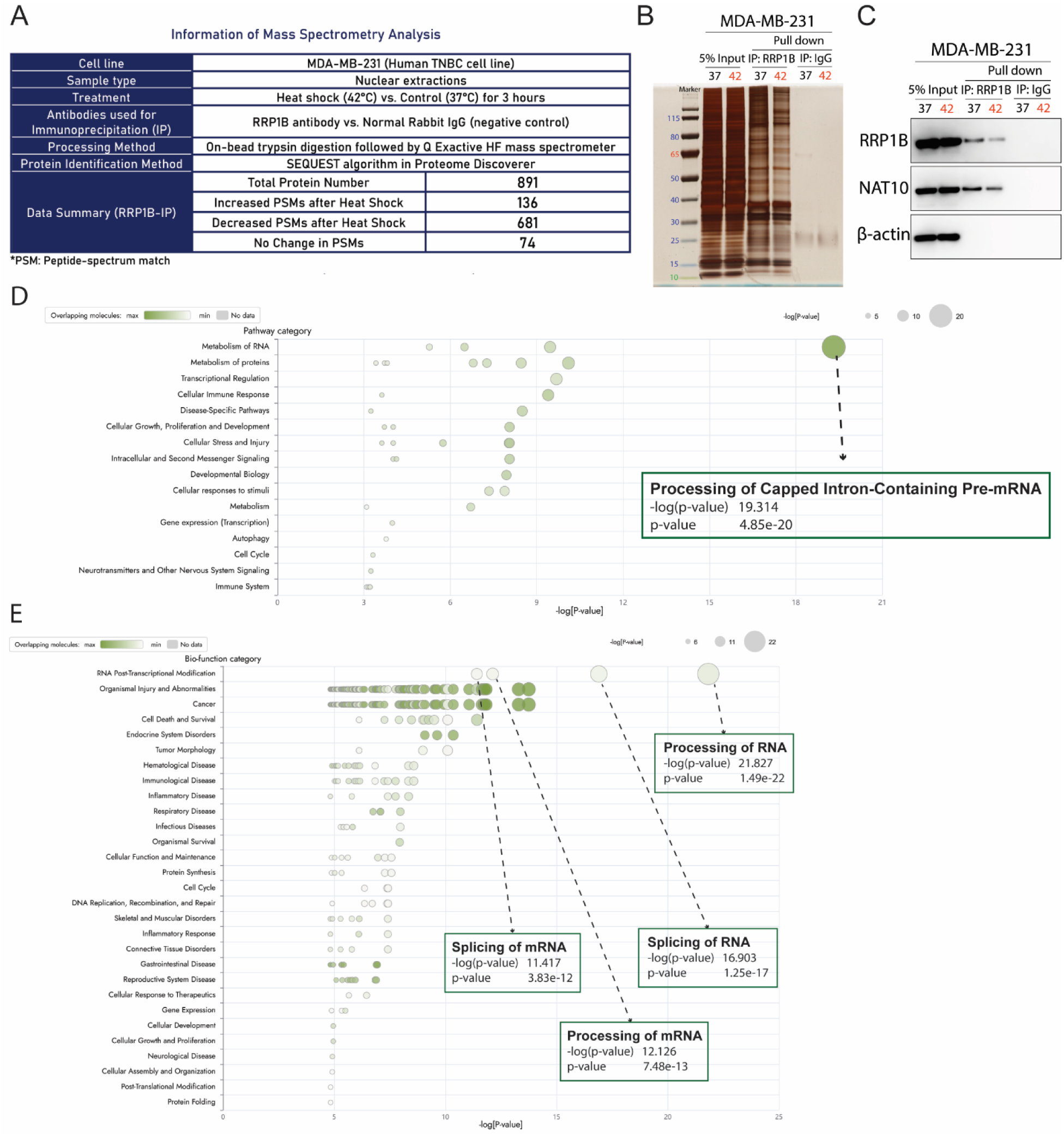
Direct association of RRP1B with RNA processing machinery in response to cellular stress, Related to Figure 3. (A) Information table summarizing the experimental design and results of the RRP1B immunoprecipitation–mass spectrometry (IP-MS) analysis. (B) Representative silver staining image of RRP1B IP from nuclear extracts of parental and RRP1B-overexpressing MDA-MB-231 cells following 3-hour treatment at 37 °C or 42 °C. (C) Representative Western blots of RRP1B IP from nuclear extracts of parental and RRP1B-overexpressing MDA-MB-231 cells following 3-hour treatment at 37 °C or 42 °C. (NAT10: a previously identified RRP1B-binding protein by our lab^69,70^) (D-E) Bubble charts of the top canonical pathways (D) and disease/function categories (E) identified by Ingenuity Pathway Analysis (IPA) among the 136 proteins with increased RRP1B binding under stress conditions.

**Figure S4.**
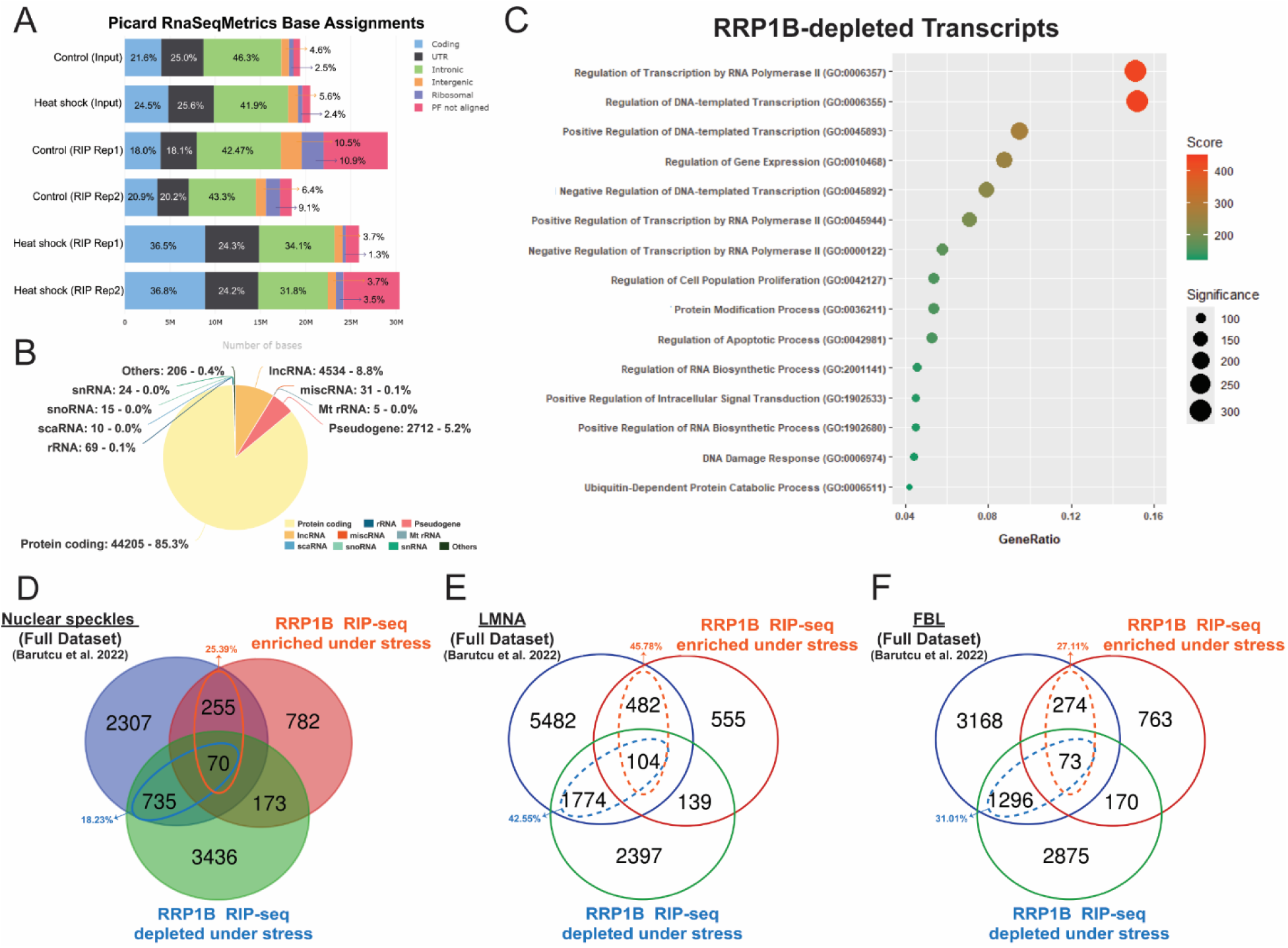
Differential RRP1B binding to RNAs derived from protein-coding regions, related to Figure 4. (A) Proportional bar chart showing the base composition of RIP-seq samples. (“PF not aligned” reads were excluded from percentage calculations.) (B) Pie chart showing the RNA biotypes and their proportion identified from RRP1B RIP-seq analysis. (C) Dot plot of the top enriched Gene Ontology pathways among RRP1B-depleted transcripts under stress conditions. (D-F) Venn diagrams showing the overlapping transcripts between APEX-seq datasets^7^ and RRP1B-bound transcripts in stress conditions. (LMNA: nuclear membrane marker; FBL: nucleolar marker)

**Figure S5.**
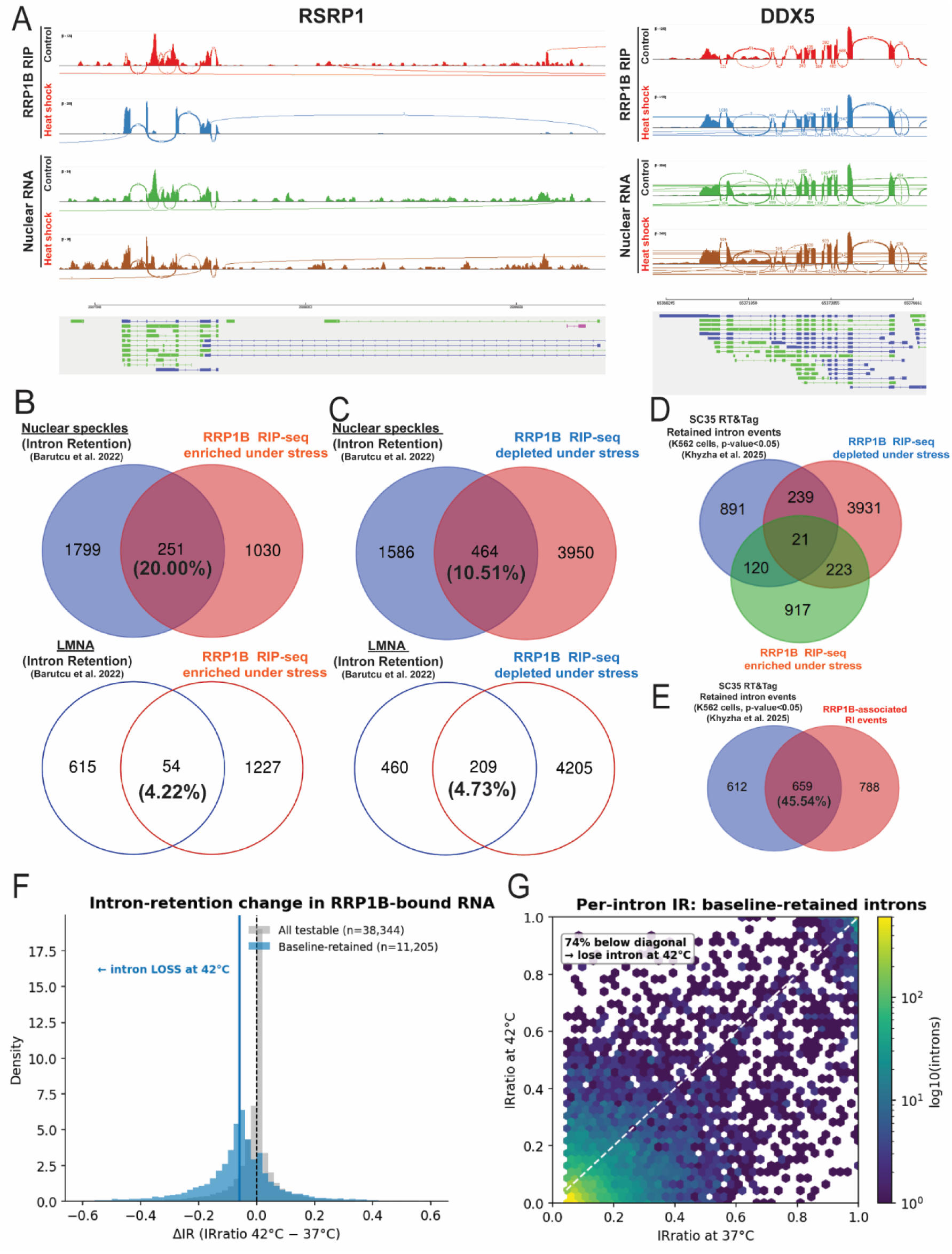
Stress-induced changes in RRP1B-associated intron retention, related to Figure 5. (A) Sashimi plots illustrating several representative transcripts with significant RRP1B-associated retained-intron (RI) splicing events. (RSRP1: Arginine and serine rich protein 1; DDX5: DEAD-Box Helicase 5) (B-C) Venn diagrams of overlapping RI events among NS APEX-seq, nuclear membrane (LMNA) APEX-seq, and stress-associated RRP1B-bound transcripts. (D) Venn diagram showing the overlapping transcripts between SC35 reverse transcribe and tagment (RT&Tag) RI dataset and RRP1B-bound transcripts in stress conditions. (E) Venn diagram showing the overlapping transcripts between SC35 reverse transcribe and tagment (RT&Tag) RI dataset and RRP1B-bound RI events in stress conditions. (F) Bar chart of intron retention ratios for RRP1B-bound and background introns at 37 °C and 42 °C. (ΔIR: changes in intron retention ratio) (G) Diagonal scatter plot of IR ratio changes in RRP1B-associated RI between 37 °C and 42 °C.

**Figure S6.**
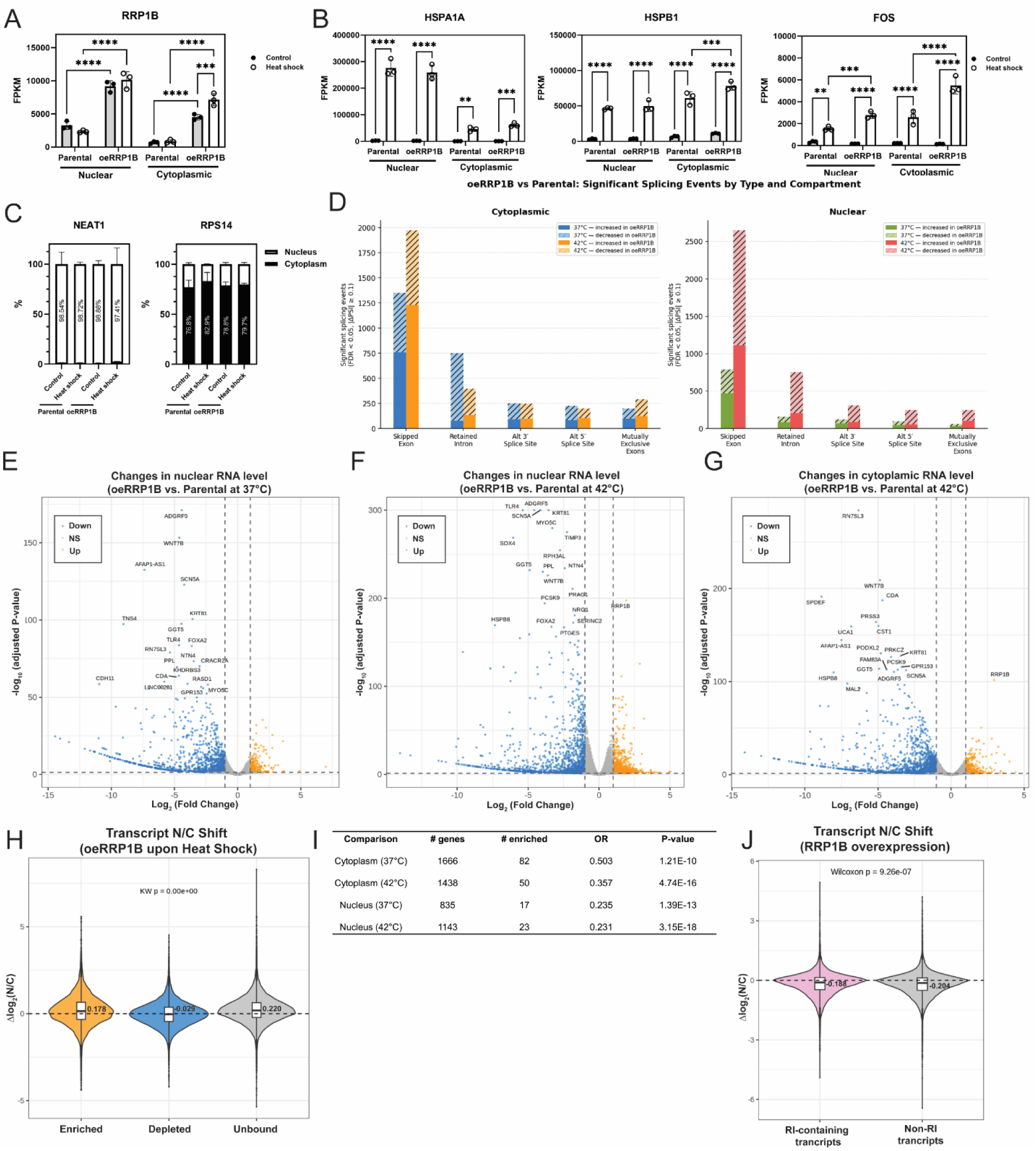
RRP1B-induced changes in subcellular transcriptomic programs, related to Figure 6. (A) Quantitative analysis of RRP1B level in subcellular fractions of MDA-MB-231 parental and RRP1B overexpressing cells treated with either 37°C or 42°C for 3 hours. (Two-way ANOVA with Fisher’s LSD test. ***, p<0.0005, ****, p<0.0001) (B) Quantitative analysis of the expression of known stress response genes, including HSPA1A, HSPB1, and FOS, in subcellular fractions of MDA-MB-231 parental and RRP1B overexpressing cells treated with either 37°C or 42°C for 3 hours. (Two-way ANOVA with Fisher’s LSD test. **, p<0.005, ***, p<0.0005, ****, p<0.0001.) (C) Quantitative analysis of NEAT1, a known nuclear-enriched RNA, and RPS14, a ribosomal RNA that is cytoplasmically enriched, in subcellular fractions of MDA-MB-231 parental and RRP1B overexpressing cells treated with either 37°C or 42°C for 3 hours. (Two-way ANOVA with Fisher’s LSD test.) (D) Bar charts showing the distribution of differential splicing events, including skipped exons (SE), retained introns (RI), alternative 3′ splice sites (A3SS), alternative 5′ splice sites (A5SS), and mutually exclusive exons (MXE), across different subcellular fractions and temperature conditions following RRP1B overexpression. (E-G) Volcano plots showing RRP1B overexpression–induced changes in nuclear RNA expression at 37 °C (E) and 42 °C (F), and cytoplasmic RNA expression at 42 °C (G). (H) Representative RNA-FISH images of MDA-MB-231 and Hs578T cells treated with heat shock (42°C) for 3 hours, acidic shock (pH6.5) for 1 hour, or under normal culture conditions (37°C, pH7.4). Scale bar: 10 μm. (I) Violin plot of transcript N/C ratio shifts in RRP1B-enriched, RRP1B-depleted, and unbound transcripts in oeRRP1B cells. (Kruskal–Wallis test.) (J) Summary table showing the representation of RRP1B-associated genes among differentially expressed genes across all compartments and temperature conditions. (K) Violin plot of transcript N/C ratio shifts in RI-containing, non-RI transcripts in oeRRP1B cells. (Wilcoxon test.)

**Figure S7.**
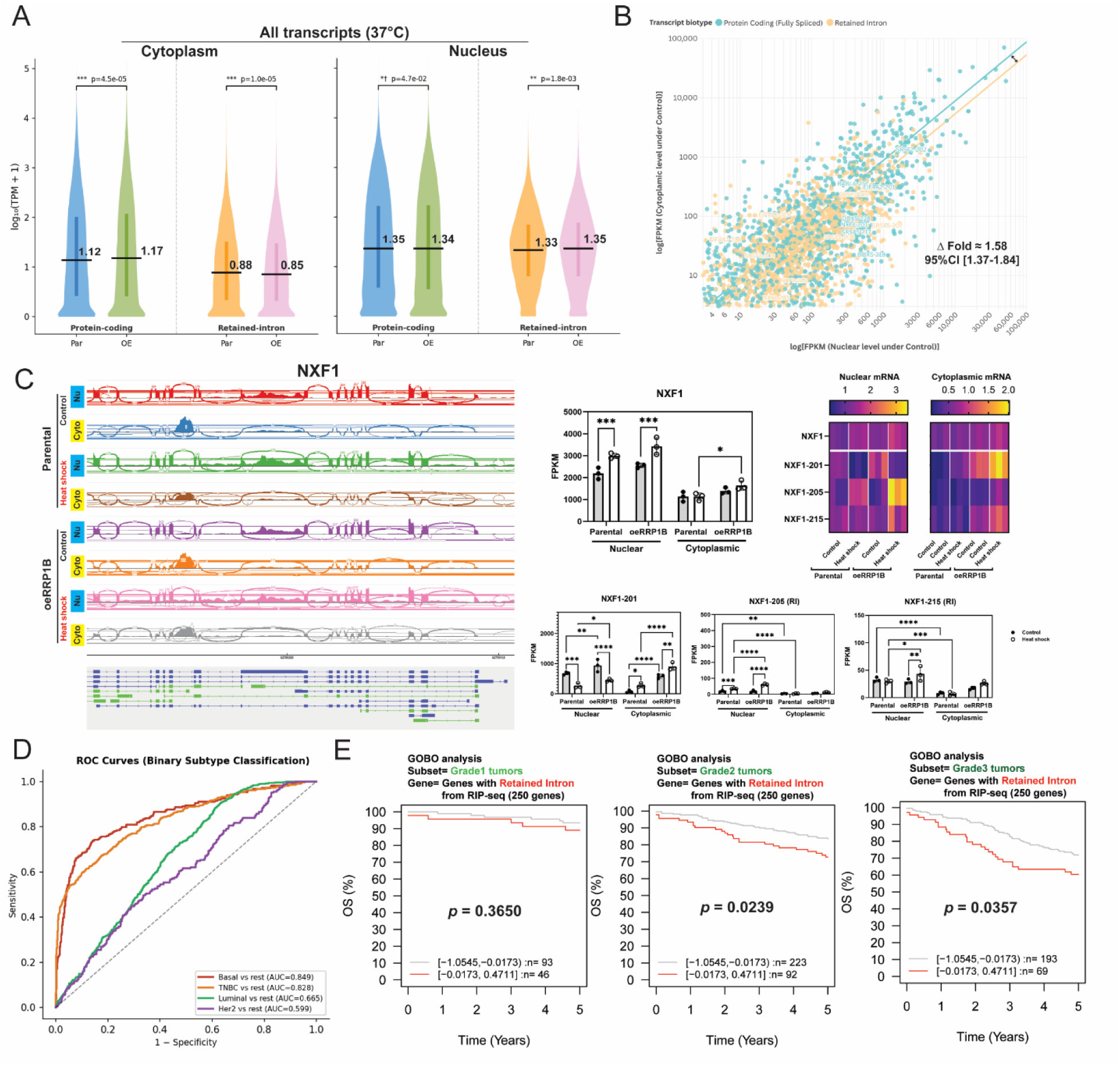
RRP1B promotes export of fully spliced transcripts from retained-intron–containing genes, related to Figure 7. (A) Violin plots comparing protein-coding and retained-intron (RI) isoforms across subcellular fractions for all genes between parental and oeRRP1B cells. (Mann-Whitney U test.) (Par: Parental, OE: RRP1B overexpression) (B) Scatter plot with trend lines showing the ratio between nuclear and cytoplasmic levels in protein-coding and RI transcripts among 250 RI genes in MDA-MB-231 parental cells under control conditions. (CI: Confidence Interval) (C) Sashimiplots, quantitative analyses, and heatmaps showing the changes in RRP1B-binding and subcellular RNA expression of NXF1 transcript isoforms upon stress. (Two-way ANOVA with Fisher’s LSD test. *, p<0.05, **, p<0.01,***, p<0.0005, ****, p<0.0001) (D) Receiver operating characteristic (ROC) curves illustrating the ability of the RRP1B-associated transcriptomic signature to distinguish individual METABRIC PAM50 subtypes. (E) GOBO analysis of the 164-gene RRP1B-associated RI signature in overall survival of breast cancer patients by tumor grade.

## Figure legends of other supplementary materials

**Table S1**. List of primers and antibodies used in this paper.

**Table S2.** List of proteins identified by RRP1B immunoprecipitation–mass spectrometry (IP-MS) analysis, related to Figure 3.

**Table S3**. List of RRP1B-enriched and RRP1B-depleted transcripts identified by RNA immunoprecipitation sequencing (RIP-seq) analysis, related to Figures 4-6.

**Table S4**. List of genes overlapping between RRP1B-associated transcripts and public datasets, related to Figures 4–5.

**Table S5**. List of transcripts identified by rMATS-based alternative splicing analysis of RRP1B RIP-seq data, related to Figures 5 and 7.

**Supplementary PRALN File.** Sequence alignment of human, chicken, and platypus RRP1B proteins.

**Movie 1.** Representative 3D immunofluorescence reconstruction generated from Z-stack images of HS578T cells expressing GFP-tagged RRP1B-S350A (unphosphorylatable mutant) (Red Fluorescence signals: blue, DAPI; green, GFP; red, SC35)

**Movie 2.** Representative 3D immunofluorescence reconstruction generated from Z-stack images of HS578T cells expressing GFP-tagged RRP1B-S350D (Phosphomimetic mutant) (Red Fluorescence signals: blue, DAPI; green, GFP; red, SC35)

